# Evaluation of false positive and false negative errors in targeted next generation sequencing

**DOI:** 10.1101/2024.07.22.603478

**Authors:** Youngbeen Moon, Young-Ho Kim, Jong-Kwang Kim, Chung Hwan Hong, Eun-Kyung Kang, Hye Won Choi, Dong-eun Lee, Tae-Min Kim, Seong Gu Heo, Namshik Han, Kyeong-Man Hong

## Abstract

**Background:** Although next generation sequencing (NGS) has been adopted as an essential diagnostic tool in various diseases, NGS errors have been the most serious problem in clinical implementation. Especially in cancers, low level mutations have not been easy to analyze, due to the contaminating normal cells and tumor heterozygosity.

**Results:** In targeted NGS (T-NGS) analyses for reference-standard samples containing mixtures of homozygote H. mole DNA with blood genomic DNA at various ratios from four certified NGS service providers, large differences in the lower detection limit of variants (16.3 times, 1.51∼24.66%) and the false positive (FP) error rate (4280 times, 5.814 x 10^−4^ ∼1.359 x 10^−7^) were found. Employment of the commercially available Dragen system for bioinformatic analyses reduced FP errors in the results from companies BB and CC, but the errors originating from the NGS raw data persisted. Bioinformatic conditional adjustment to increase sensitivity (less than 2 times) led to a much higher FP error rate (610∼8200 times). In addition, problems such as biased preferential reference base calls during bioinformatic analysis and high-rate FN errors in HLA regions were found in the NGS analysis.

**Conclusion:** T-NGS results from certified NGS service providers can be quite various in their sensitivity and FP error rate, suggesting the necessity of further quality controls for clinical implementation of T-NGS. The present study also suggests that mixtures of homozygote and heterozygote DNAs can be easily employed as excellent reference-standard materials for quality control of T-NGS.

## 1 Introduction

Next-generation sequencing (NGS) technology can examine millions of DNA variants at a time, and over 11,000 laboratories and companies in the USA have developed their own NGS tests for genetic diagnosis with a revenue of 8.8 billion in 2020 [20, 57] (https://www.ncbi.nlm.nih.gov/gtr/), and enabling discoveries of a lot of cancer-associated or disease-causing mutations [44, 8, 42]. Despite widespread adoption of NGS technology for clinical diagnosis, high error rates on various NGS platforms have been reported (0.26-12.86%) [50]. In a study with 20,000 samples, a NGS panel test showed false positive (FP) error rate of 1.3%, and the authors recommended confirmation with Sanger sequencing [43]. We also reported that whole exome NGS results had high rate (40-45%) of false negative (FN) errors by performing targeted NGS (T-NGS) analysis in cancer cell lines, and the high FN errors are related to most of the 43% mutation call inconsistencies between GDSC (Genomics of Drug Sensitivity in Cancer) and CCLE (Cancer Cell Line Encyclopedia) cell line mutation databases [30], indicating that low sensitivity of whole exome NGS analyses is making FN errors. These reports suggest the necessity for the evaluation of NGS sensitivity and specificity.

The Food and Drug Administration (FDA) reported a guidance document to accelerate the establishment of a regulatory approach for NGS testing [15, 37]. Currently, the clinical laboratories for NGS testing are overseen by the Centers for Medicare and Medicaid Services (CMS) according to the Clinical Laboratory Improvement Amendments (CLIA) regulations [51] in the USA. A lot of studies on the quality control of NGS technologies have been published via SEQC2 (Sequencing Quality Control 2) project or by independent researchers including the best practice in cancer mutation detection with whole-genome and whole-exome sequencing [66, 70]; establishment of reference samples for sequencing [13, 26, 40] or for single-cell RNA sequencing technology [5, 45, 12]; and performance assessment of various DNA sequencing platforms [16]. These studies indicate the necessity of further NGS validation technologies and of more independent reference-standard materials.

T-NGS has been adopted for clinical testing to detect mutations in tumor samples which can give a guide to select the best anti-cancer agents [29, 34, 33, 27, 59, 24]. However, detection of mutations for tumor samples is a lot more difficult than that for genetic diseases with NGS technology, due to genetic heterogeneity of cancer cells even in a tumor mass and the presence of contaminating normal cells [14, 53, 67, 1]. Therefore, the evaluation of lower detection limit and their specificity of mutation calls will be essential for T-NGS testing for tumor samples, and the mixture of many cell lines or the employment of spiked mutation sequences have been suggested as reference standards for T-NGS quality control [17, 56, 41, 54]. Studies via SEQC2 compared the sensitivity and accuracy of T-NGS for eight Pan-Cancer panels [19, 18]using reference standards, however, for broader applications of the reference standards at each laboratory and clinic, still further investigation is needed: the employment of mixture of many cell lines or DNAs as a reference standard necessitates quantitative pre-validation of each variant alleles using whole genome/exome sequencing and droplet digital PCR methods [26], and the presence of minor variant alleles with low level allelic fraction (AF) within each cancer cell line [30] can interfere precise estimation of T-NGS specificity. In the present study, we developed a new method for the evaluation of NGS kits or services without pre-characterization studies on the reference standard, and this can be applied to almost all targeted NGS or whole exome NGS kits from various technologies. To validate T-NGS kits, in the present study, the NGS service providers which are accredited by College of American Pathologists (CAP) or Ministry of Food and Drug Safety (Korean FDA) for NGS testing were chosen, and we evaluated the lower detection limits and false positive error rates from four T-NGS results for the reference-standard DNAs.

## 2 Results

### Evaluation of T-NGS sensitivity from the results by inhouse bioinformatic analysis for reference-standard DNAs

For the evaluation of T-NGS sensitivity from various NGS service providers, reference-standard DNAs were prepared: DNA1 (which is H. mole DNA and has only homozygote alleles) and DNA2 (which is blood DNA from an individual) were mixed at various ratios, and the mixed DNA samples were employed as reference standards for T-NGS analysis. In the analysis of sensitivity, informative alleles are the alleles in which variant or reference base is detected in either DNA1 or DNA2 but not both. There are 11 informative alleles for sensitivity in an example of Fig. 1A. Three of the alleles are the null and homozygote pair alleles (N-Ho pairs), which are null in DNA1 and homozygote variant in DNA2; three of the alleles are homozygote and null pair alleles (Ho-N pairs) which are homozygote variants in DNA1 and nulls in DNA2; and 5 of the alleles are null and heterozygote pair alleles (N-He pairs), which are null in DNA1 and heterozygote variant in DNA2. Sensitivity of T-NGS is determined by the least percentage of allele fraction at which 95% of the diluted informative alleles are detected, and the sensitivity of NGS in the example of Figure 1A is more than 2.5% AF because none of N-He pair alleles were detected in samples of CH5 (DNA1: DNA2 = 5:95) which has informative variant alleles with 2.5% AF, suggesting that no alleles with 2.5% AF are detected. In contrast, the N-Ho pair alleles for CH95 (DNA1: DNA2 = 95:5), Ho-N pair alleles for CH5 (DNA1:DNA2 = 5:95), and N-He pair alleles for CH90 (DNA1: DNA2 = 80:20), all of which has variant alleles with 5% AF, are detected in the DNA mixture samples, suggesting that T-NGS sensitivity is between 2.5 and 5% in AF.

**Figure 1:**
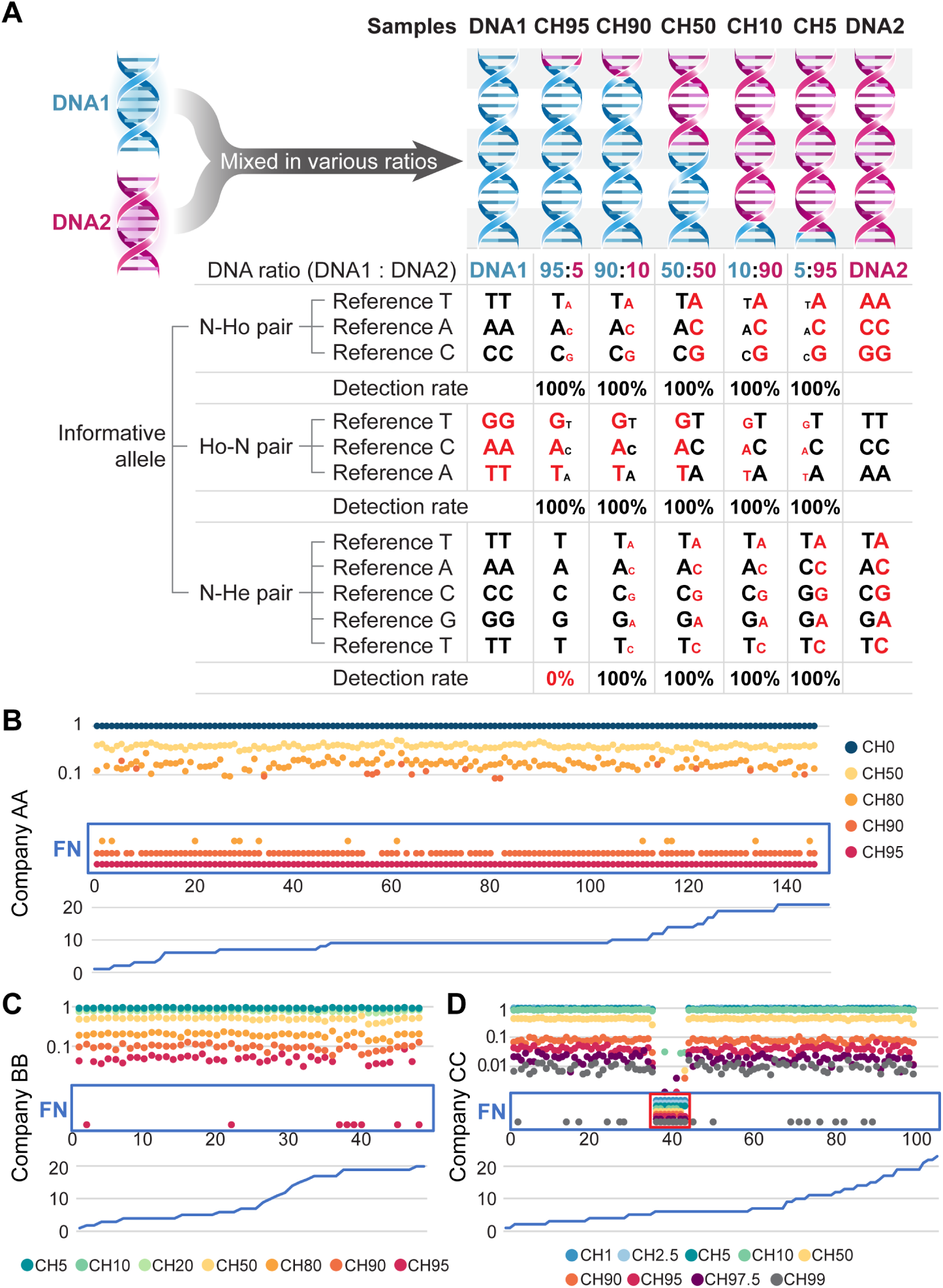
Evaluation of T-NGS sensitivity by estimating FN errors in mixed reference standards. A. Sensitivity determination with T-NGS results from reference standards. Evaluation of sensitivity in NGS results employing the reference-standard DNAs which were prepared by mixing DNA1 (H. mole DNA) and DNA2 (blood DNA) at various ratios There are three types of informative alleles such as N-Ho pair, Ho-N pair, and Ho-He pair. The sensitivity of the NGS results is between 2.5 and 5% in the example. Variant alleles detected were marked in red, and the relative ratio of reference and variant bases were marked by the relative size of characters. B. FN errors in the company AA for N-Ho pair alleles. All 5% and most of 10% informative alleles were not detected, suggesting that the sensitivity was about 20%. C. FN errors in company BB for N-Ho pair alleles. Most of informative alleles with 5% or more expected allelic fraction (AF) were detected. D. FN errors in company CC for N-Ho pair alleles. Most 1% informative alleles were detected, and the sensitivity was about 1%. FN alleles at mixed reference standards were shown within the red rectangle. At the lower columns, chromosome numbers were shown. The ratios of DNA1 and DNA2 for each sample (CH0 ∼CH100) were shown in Materials and Methods. X-axis, alleles aligned by chromosome and position. Y-axis, VAF.

T-NGS experiments and bioinformatic analysis were performed by 3 companies (AA to CC) for the reference-standard DNAs in addition to the unmixed samples of DNA1 and DNA2. Among variant calls from T-NGS results for reference-standard samples, N-Ho pair informative alleles were analyzed first. From the N-Ho pair alleles by each company, VAF values for the mixed DNA samples were aligned by chromosomal positions as shown in Figure 1. After the evaluation of the positive and FN alleles, the least percentage of allele fraction at which 95% of the diluted informative alleles were about 20% and 5% for the T-NGS results from companies AA and BB (Figs. 1B and 1C).

At some alleles in the results from company CC, there were FN alleles in all mixed DNA (or variant-diluted) samples, resulting in no variant-dilution effect (marked in green box in Fig. 1D), which is a linear relationship between expected VAF values and the VAF values from T-NGS analyses. All FN error alleles showing no variant-dilution effect were found to be in HLA region of chromosome 6, suggesting that the variant calls for alleles in HLA region may not be accurate during bioinformatic analysis, and led to FN error calls. When the alleles in chromosome 6 were removed, the sensitivity of T-NGS results from company CC was about 1%. Our results indicate that the variation of T-NGS sensitivities from various companies could be as large as 20 times (1% to 20%), which is quite comparable to the result (1 ∼5%) from a recent report by SEQC consortium [19], except the alleles in HLA region at which the dilution effects were not observed.

The results for Ho-N pair alleles from companies BB and CC were analyzed as in Fig. S1. Ho-N pair alleles from company AA were not analyzed due to no analysis of reference standards for low level diluted variants in DNA1, for example CH5, CH10, and CH20. The sensitivities of T-NGS results from company BB and CC for Ho-N pair alleles were about the same as those for N-Ho pair alleles. In addition, the Ho-N pair alleles in the HLA regions (chromosome 6) showed no variant-dilution effect.

In the analysis of the sensitivity in He-N pair alleles from T-NGS results of company AA, most variant alleles with 20% AF were not detected but most variant alleles with 50% AF were detected (Fig. S2A), suggesting that the sensitivity was about 25% which is consistent with that from Ho-N pair alleles. Most of He-N alleles with 5% VAF were not detected in the results from company BB (Fig. S2B), suggesting that the sensitivity of T-NGS of company BB was more than 2.5%. Most of He-N pair alleles with 1% VAF were not detected in the results from company CC (Fig. S2C), suggesting that the sensitivity of T-NGS of company CC was higher than 0.5%. These results from companies BB and CC are also consistent with those for N-Ho or Ho-N pair results from companies BB and CC, respectively.

### Correlation skewing between expected VAFs and VAFs from T-NGS results and its correction by Dragen system

Reliability of VAF values from T-NGS results can be evaluated by the correlation with the expected VAFs which can be calculated by the dilution folds of the variants in the mixed reference-standard samples [36]. In the analyses of the correlation in three companies, overall correlation coefficients were over 0.99 for companies BB, CC, and DD (Fig. 2), but the correlation for company AA was skewed with the correlation coefficient < 0.98 (Fig. 2A): the skewing in company AA was apparent epecially in the alleles with high dilution folds (lower part of the line).

**Figure 2:**
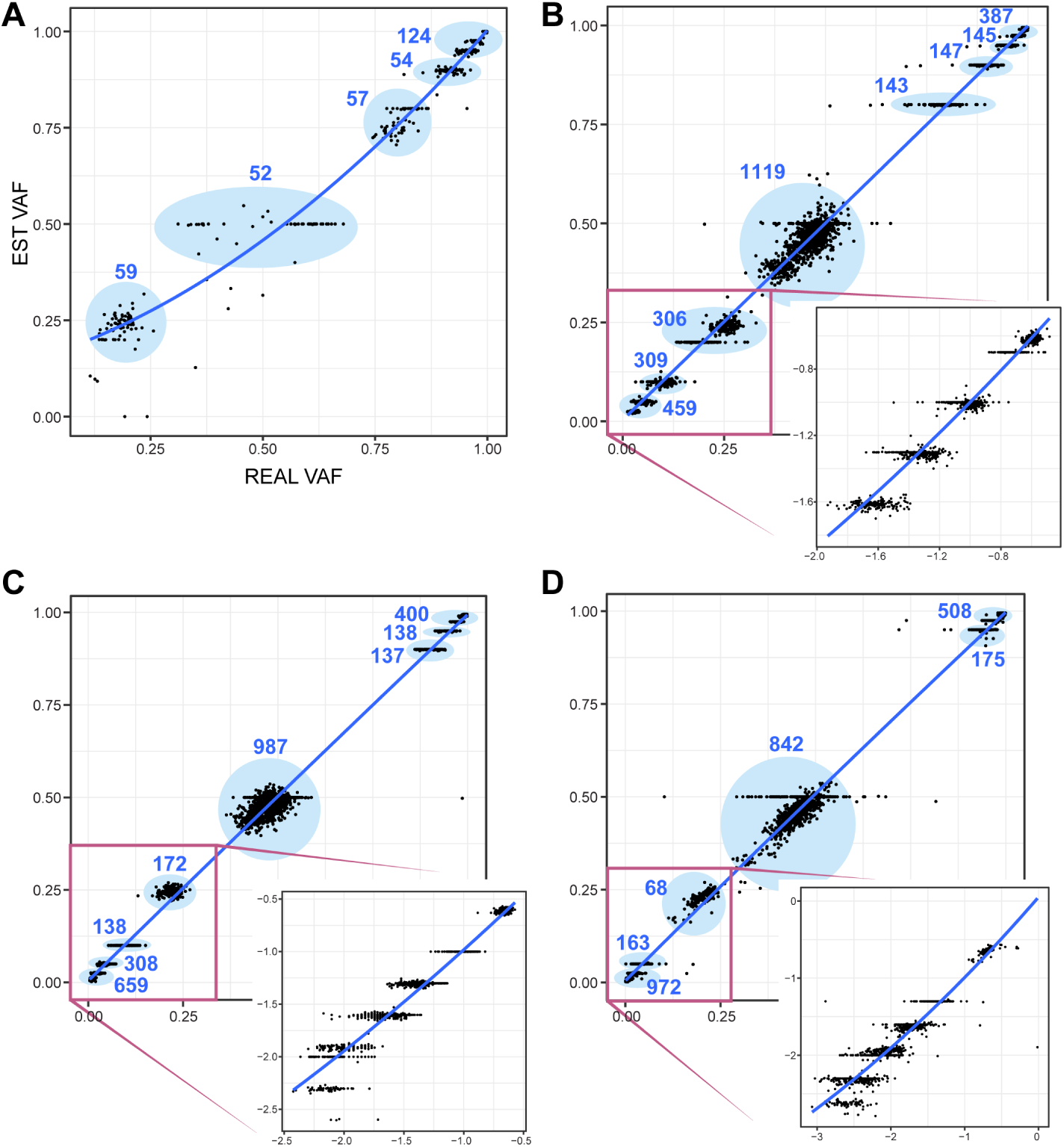
Correlation of allelic fraction value from T-NGS analysis with the expected allelic fraction value calculated based on dilution folds of homozygote or heterozygote variants. A. The correlation of allelic fractions for variants by company AA inhouse method (AA-inhouse, R = 0.9789). B. The correlation of allelic fractions for variants by company BB inhouse method (BB-inhouse, R = 0.9969). C. The correlation of allelic fractions for variants by company CC inhouse method (CC-inhouse, R = 0.9935). D. The correlation of allelic fractions for variants by company DD inhouse method (DD, R = 0.9964). The numbers of informative alleles in the circle were shown. For figures B to D, a part of lower left area (marked as a rectangle) was re-drawn at the lower right corner with the scale modification. X-axis, VAF values from T-NGS analysis. Y-axis, the expected VAF values.

The reason for the correlation skewing may be either from the raw data generation or from bioinformatic analytic process. To dissect the origin of the correlation skewing, the whole raw data were analyzed by the bioinformatic analysis software of Dragen system (version 4.2, Illumina Korea, Seoul). For Dragen analysis, bed file is needed, but the companies of AA and CC refused to provide the bed files, and therefore only exons of target genes were anlyzed by Dragen system for the fastq files from companies AA and CC. In the bioinformatic analysis result by Dragen for company AA, the skewing is much less evident, and the correlation was more linear (R = 0.9856, Fig. S3), suggesting that the AA-inhouse method for bioinformatic analysis was the major reason for the skewing, indicating that inhouse bioinformatic analysis of company AA may skew the results of T-NGS.

The T-NGS system for three companies did not use unique molecular identifier (UMIs) system for high sensitivity and specificity of T-NGS results [32, 63, 60]. To test the effect of UMI on T-NGS analyses, we sent the same reference-standard samples to company CC, which generated raw data with a T-NGS kit adopting UMIs. Variant analysis from T-NGS raw data adpoting UMIs was performed by the provider’s bioinformatic tools (inhouse method), and the analytic results are marked as DD in Fig.2D.

### Comparison of T-NGS sensitivity analyzed by either inhouse or Dragen bioinformatic system

After N-Ho pair, Ho-N pair and He-N pair data were merged by the expected dilution folds of the variants, all sensitivity data from 4 companies (AA to DD) were plotted in Fig. 3. The inhouse results from company DD was modified again by selecting variant alleles only with VAF *≥* 0.005, and the sensitivity result was marked as DD AF *≥* 0.005 (Fig. 3). The alleles on chromosome 6 were not included for company CC, because VAF of the alleles did not show variant-dilution effects as shown in Fig. 1D, Additional file 1: Fig. S1B, and Fig. S2C. The least percentage of VAF at which 95% of the diluted informative alleles are detected were from 1.51% (company DD inhouse method) to 24.7% (company AA dragen method): the sensitivity difference was 16.3 times (Fig. 3). We also found that the sensitivities from Dragen analysis for companies AA, BB, and CC were similar to those from inhouse methods, and the difference was much less than 2 folds (0.849∼1.140).

**Figure 3:**
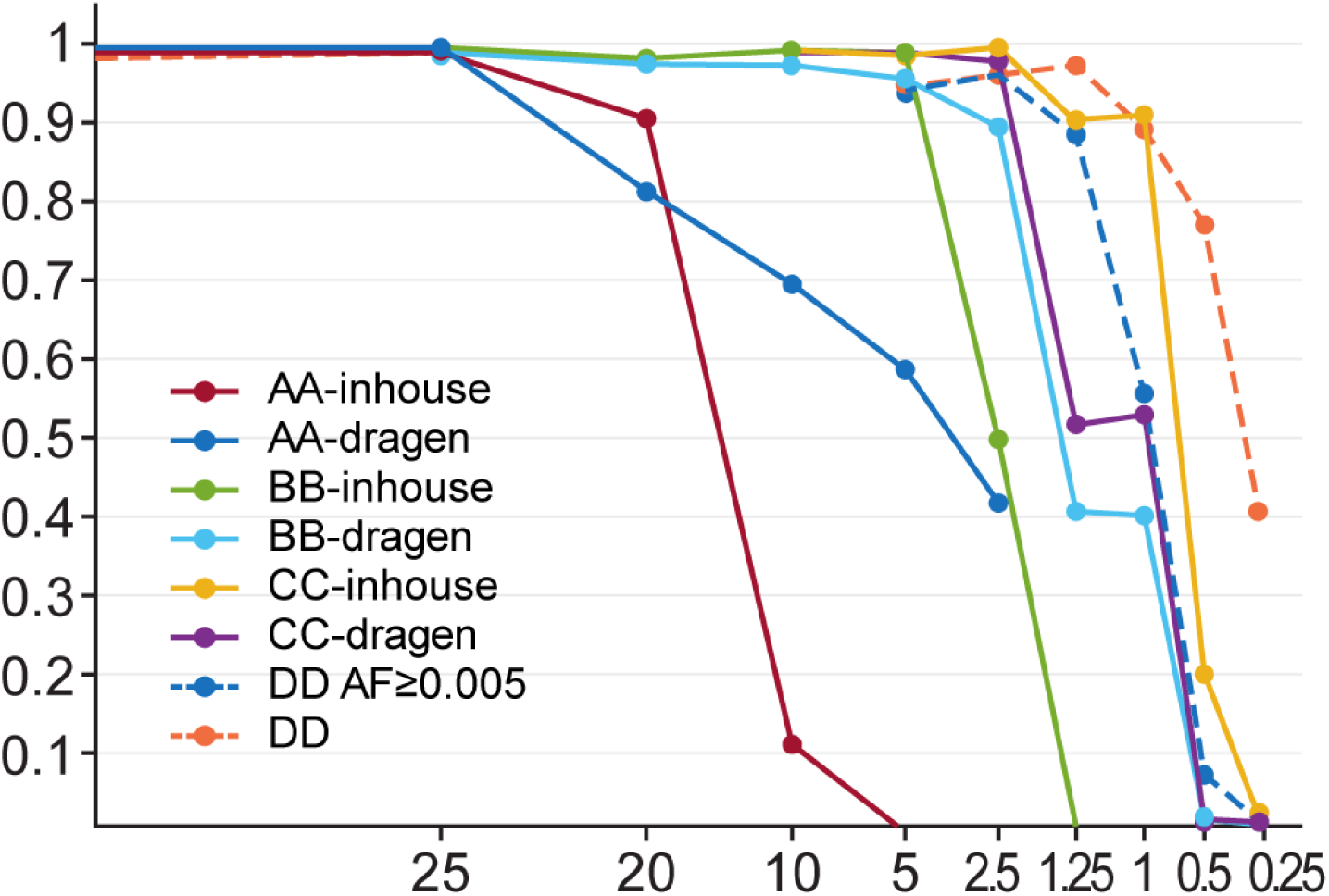
Sensitivity of various T-NGS methods depending on companies and analytic methods. Inhouse bioinformatic analysis was performed by each company’s own bioinformatics methods. For the comparison of bioinformatic performance, variant analyses were also performed by Dragen system on the fastq files from each company. X axis is the expected percentage of VAF for the informative alleles, and Y-axis is the detection rate of informative alleles at the expected percentage of VAF. AA-dragen (least VAF for 95% detection, 24.66%), T-NGS raw data was obtained from AA, and variant call was analyzed by Dragen system; AA-inhouse (23.03%), T-NGS raw data from AA, and variant call by AA inhouse method; BB-dragen (3.99%), T-NGS raw data from BB, and variant call by Dragen system; BB-inhouse (4.55%), T-NGS raw data from BB, and variant call by BB inhouse method; CC-dragen (2.25%), T-NGS raw data from CC, and variant call by Dragen system; CC-inhouse (1.91%), T-NGS raw data from CC, and variant call by CC inhouse method; DD AF *≥* 0.005 (1.98%), variants with AF *≥* 0.005 from the variant calls by DD inhouse method; DD (1.51%), variant call by DD inhouse method for T-NGS raw data from DD.

The sensitivity of T-NGS is usually related to sequencing depth and quality of generated raw data [25, 47, 11]. The sequencing depths of the results from each company were plotted along with chromosomal sites, and it was expected that T-NGS sensitivity was closely related to sequencing depth (Fig. S4). However, the linear relationship between the sensitivity and the sequencing depth was not perfectly matched: 1) although the T-NGS results from companies CC and DD showed similar sensitivity (Fig. 3), the sequencing depth for DD was high than that for CC (about 1.91-fold difference). 2) the sequencing depth of the results from companies BB and CC were similar (less than 1.07-fold difference), but the sensitivity difference was about 2 times higher in CC. These results suggest that the sequencing depth is not the sole factor for the sensitivity of NGS.

Another factor for the T-NGS sensitivity may be the quality of NGS results, and per base sequence quality is usually employed for the quality check [25, 35, 55]. Representative per base sequence quality results were shown in Fig. S5. Except for a little drop of quality in the results from company AA, no significant quality problem was found, suggesting that the correlation with sensitivity may not be inferred from the per base sequence quality. Therefore, there should be a lot more factors determining T-NGS sensitivity, including the efficiency of target gene capture and the efficiency of the captured fragment amplification, which cannot be inferred from the bioinformatic parameters. Therefore, we need more direct quantitative assessments of T-NGS sensitivity as in the present study.

### Evaluation of T-NGS specificity using reference standards

Specificity of NGS methods for the tumor samples containing alleles with low level VAF, has not been evaluated routinely [19, 39, 3, 52], due to the limitations in practical evaluation methods. With the employment of reference-standard DNAs, in the present study, however, false positive (FP) errors can be precisely inferred. In the present study, FP error alleles were defined as the appearance of base(s) in the mixed reference-standard DNAs, but the base(s) was not shown in either DNA1 or DNA2. FP errors can occur in three pair alleles such as 1) R-R pair alleles at which both DNA1 and DNA2 have reference base; 2) V-V pair alleles at which neither DNA1 nor DNA2 has reference base; and 3) R-V pair alleles at which DNA1 and DNA2 have both reference and variant bases. In the example of Figure 4A, there are 1 FP error out of 10 R-R pair alleles, 2 in 10 V-V pair alleles, 1 in 10 V-R pair alleles. Therefore, total FP error rate is 4/30 = 0.13.

**Figure 4:**
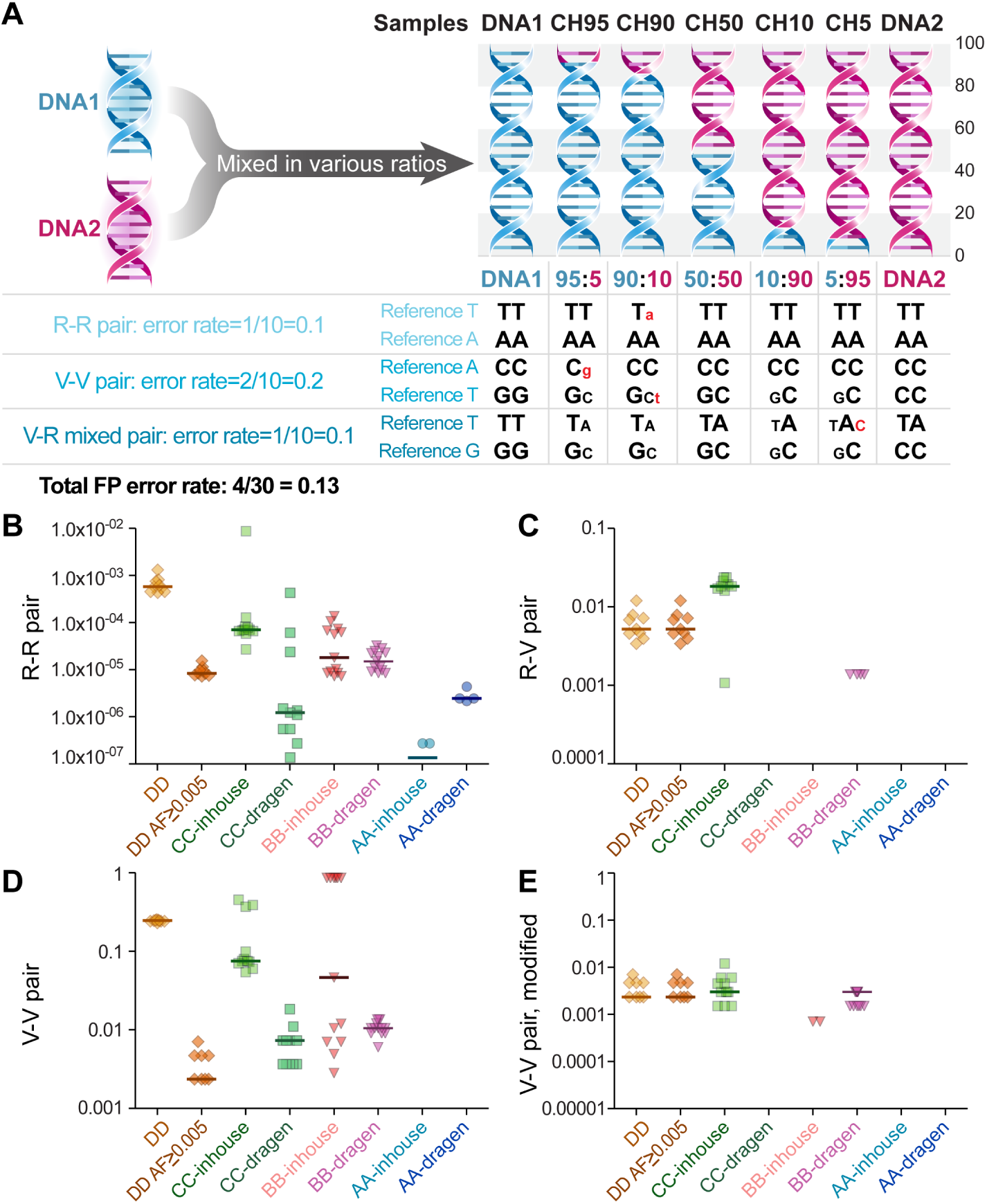
T-NGS specificity by FP error rates. A. Specificity determination based on the FP error rates of T-NGS from various companies and methods employing the reference-standard DNAs. In the example, there are 1 FP error out of 10 R-R pair alleles (both DNA1 and DNA2 have reference base), 2 out of 10 V-V pair alleles (neither DNA1 nor DNA2 has reference base), 1 in R-V pair alleles (DNA1 and DNA2 have both reference and variant bases), and total FP error rate is 4/30 = 0.13. FP error bases are marked in red. B. FP error rates in R-R pair alleles. C. False positive rates in V-R pair alleles. D. False positive rates in V-V pairs. E. FP error rates in V-V pair alleles with the removal of VVR errors. Y-axis, FP error rate. AA-inhouse, T-NGS raw data from company AA, and variant call by AA inhouse method; AA-dragen, T-NGS raw data from AA, and variant call by Dragen system; BB-dragen, T-NGS raw data from company BB, and variant call by Dragen system; BB-inhouse, T-NGS raw data from company BB, and variant call by BB inhouse method; CC-dragen, T-NGS raw data from company CC, and variant call by Dragen system; CC-inhouse, T-NGS raw data from company CC, and variant call by CC inhouse method; DD AF *geq* 0.005, T-NGS raw data from company DD, and selection of variants with AF *>* 0.005 from the variant calls by DD inhouse method; DD, T-NGS raw data from company DD, and variant call by DD inhouse method.

In the FP error rate analysis, the rate was various in R-R pairs: the lowest (1.359 x 10^−7^) in the results from company AA with inhouse method, and the highest (5.765 x 10^−4^) in the results from company DD with inhouse method. In R-V pairs for AA-inhouse (5.199 x 10^−3^), AA-dragen (5.199 x 10^−3^), and CC-inhouse (1.824 x 10^−2^), the FP error rates were higher than those for R-R pairs as shown in Figs. 4B and 4C). However, the FP error rates were extremely high (even > 0.01 in 4 methods) in most of V-V pairs (Fig. 4D). Although the number of R-R pair allele is bigger than that of R-V or V-V pair (850 times higher in company DD and 45000 times in company AA), the FP error rate in V-V pair was much higher (5954 times higher in CC-dragen or 2560 times higher in BB-inhouse) than R-R pairs (Figs. 4B and 4D). Therefore, ignorance of errors in V-V pair may skew the final FP error rates in the results from various companies.

### FP errors by the appearance of reference base at the V-V pair alleles: VVR errors

To identify the reasons for the high FP error rates in V-V pair alleles, the base changes were checked, and it was found that the FP errors occurred frequently by erroneous appearance of reference base in the mixed DNA samples at the V-V pair alleles. FP errors by erroneous appearance of reference base at the V-V pair, in the present study, were named as VVR errors. With the removal of VVR errors, it was found that the FP error rates dramatically decreased in most methods (Fig. 4E). The overall total FP error rate also decreased to a half by the removal of VVR errors (1.419 x 10^−4^ with VVR to 6.195 x 10^−5^ without VVR) in the results from company BB (Fig. S6), suggesting that the removal of VVR errors can change the total FP error rate significantly.

VVR errors may be the artifacts incurred just from bioinformatic analysis: the reference base call may be preferred during the variant calling process even for the ones with small allelic fractions, because reference base can be inferred from the reference sequence. In contrast, for the variant alleles with the same amount of allelic fraction, variant calls may not be made, because variant base cannot be inferred from the reference sequence and should be selected based on the bioinformatic conditions, resulting in a biased preferential reference base call. To test this possibility, the sensitivity employing the diluted reference bases, instead of the diluted variant bases, was analyzed from the mixed DNA reference-standard samples. As shown in Fig. S7, detection rates of reference bases were over 90% even in alleles with 0.5% reference base/variant ratio by most methods except CC-inhouse, suggesting the presence of the biased preferential reference base call during bioinformatic analysis. This result indicates that VVR errors should be removed because those are false errors originated just from the preferential reference base call not from the real FP errors during T-NGS analyses.

### FP error rate after removal of VVR errors

After the removal of VVR errors, total FP error rates were estimated (Fig. 5). The results from company AA showed the lowest FP error rate (median, 1.359 x 10^−7^). The error rate of the results from company CC with variant call by Dragen system was 1.230 x 10^−6^ (median) which was lower than inhouse results from company BB or CC. The result from company AA with variant call by inhouse method was the worst (median, 5.814 x 10^−4^). Therefore, the difference of FP error rates in various methods was as much as 4280 times (5.814 x 10^−4^/1.359 x 10^−7^). The FP error rates from the present study (5.814 x 10^−4^ ∼1.359 x 10^−7^) were similar to those by the SEQC consortium (10^−5^ ∼10^−6^ with the AF cutoff of 1%) [35], if the inhouse results from company AA (1.359 x 10^−7^) and DD (5.814 x 10^−4^) are not included due to the sensitivity issue and inclusion of alleles with VAF cutoff less than 0.5%, respectively.

**Figure 5:**
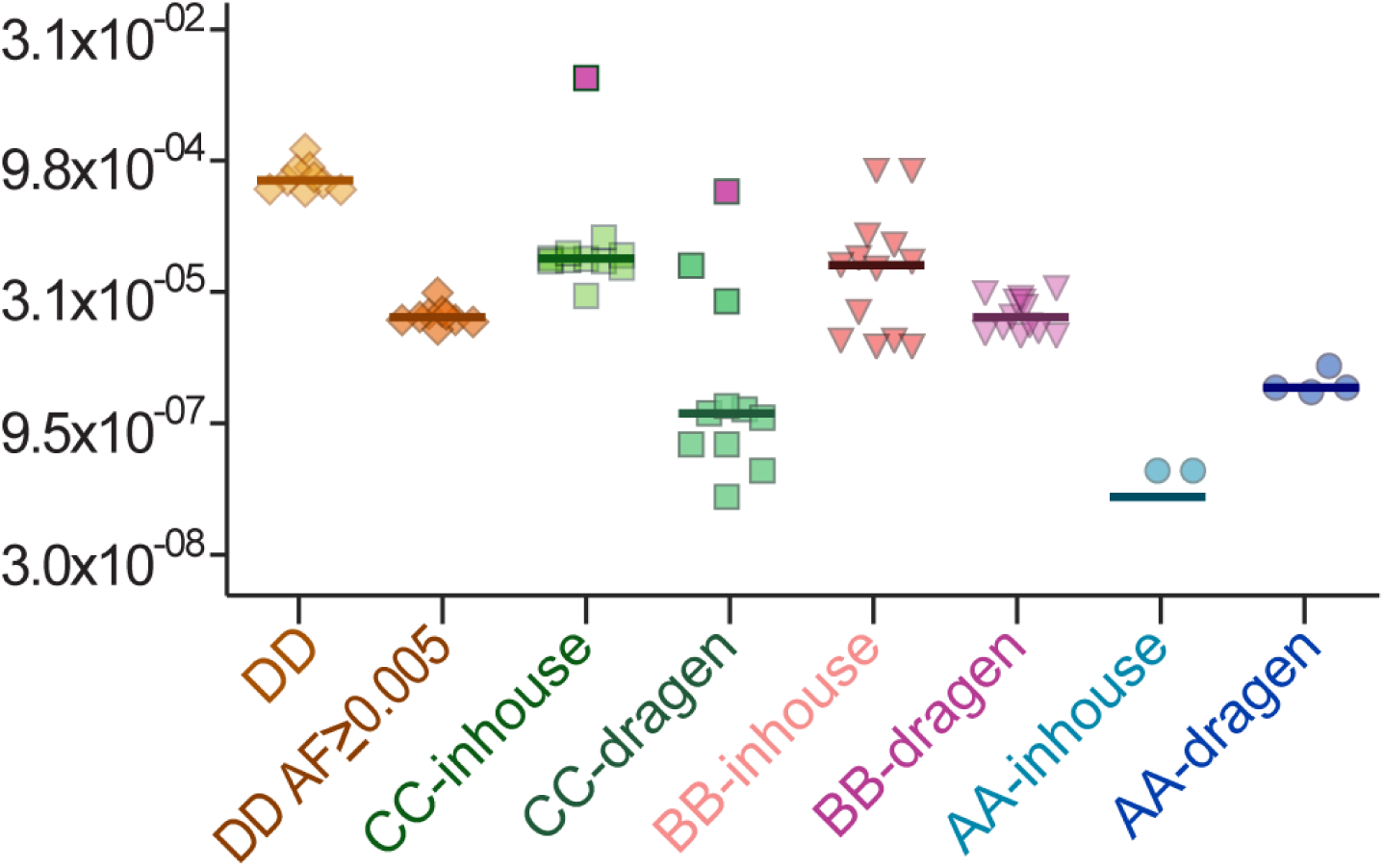
Total false positive (FP) error rates after the removal of VVR errors. Y-axis, total FP error rate. AA-inhouse (median: 1.359 x 10^−7^), T-NGS raw data from AA, and variant call by AA inhouse method; AA-dragen (median: 2.446 x 10^−6^), T-NGS raw data from AA, and variant call by Dragen system; BB-dragen (median: 1.564 x 10^−5^), T-NGS raw data from BB, and variant call by Dragen system; BB-inhouse (median: 6.195 x 10^−5^), T-NGS raw data from BB, and variant call by BB inhouse method; CC-dragen (median: 1.230 x 10^−6^), T-NGS raw data from CC, and variant call by Dragen system; CC-inhouse (median: 7.327 x 10^−5^), T-NGS raw data from CC, and variant call by CC inhouse method; DD AF *geq* 0.005 (median: 1.583 x 10^−5^), T-NGS raw data from DD, and selection of variants with AF *geq* 0.005 from the variant calls by DD inhouse method; DD (median: 5.814 x 10^−4^), T-NGS raw data from DD, and variant call by DD inhouse method. The purple squares in CC-inhouse and CC-dragen were analyzed from the same raw data (CH10) produced by company CC, and their extremely high false positivity may be originated from index hoping.

The effect of the employment of Dragen system on the FP error rate was quite various depending on the companies. The error rates significantly decreased with the employment of Dragen system for CC (0.0168 times, 1.230 x 10^−6^/ 7.327 x 10^−5^; P = 0.0016 by Mann-Whitney test), increased for AA (18.000 times, 2.446 x 10^−6^/1.359 x 10^−7^; P = 0.0294 by Mann-Whitney test). The FP error was also reduced (0.252 times) by Dragen system for company BB, but not significant (P = 0.182 by Mann-Whitney test). The error rate decreased also with the selection of alleles with AF *≥* 0.005 in the results from company DD (0.0272 times, 1.583 x 10^−5^/ 5.814 x 10^−4^; P = 0.0001 by Mann-Whitney test). These results suggest that the effects of the employment of Dragen system on the FP error rates are largely depending on variant analysis conditions for the inhouse methods. For example, the variant call conditions in inhouse methods for company CC might be looser than Dragen default condition, resulting in more FP errors. In contrast, the condition in inhouse method for company AA might be tighter than Dragen default condition for company AA, resulting in less FP errors.

### FP error-prone alleles

To infer the reasons for the FP errors, the positional information of the FP alleles was analyzed for the results from company CC (Fig. 6). At a lot of FP error alleles (37.8%, 801/2117), more than one errors were found in the mixed reference standards, and therefore as few as 9.92% FP alleles (N = 210) comprised of 32.4% of total FP errors (N = 1419). These results suggest that the presence of FP error-prone allelic sites, and the removal of these sites from the analysis may reduce FP errors. However, the distribution of FP error alleles changed with their second batch service for CH0.5 and CH99.5 samples. In addition, FP error alleles in the CH95 results which was done by the first batch service are quite similar to the results from 2nd batch service, suggesting that the FP errors even at the error-prone alleles are not stable and that the errors can be reduced just by the changes in experimental conditions for raw data production of T-NGS.

**Figure 6:**
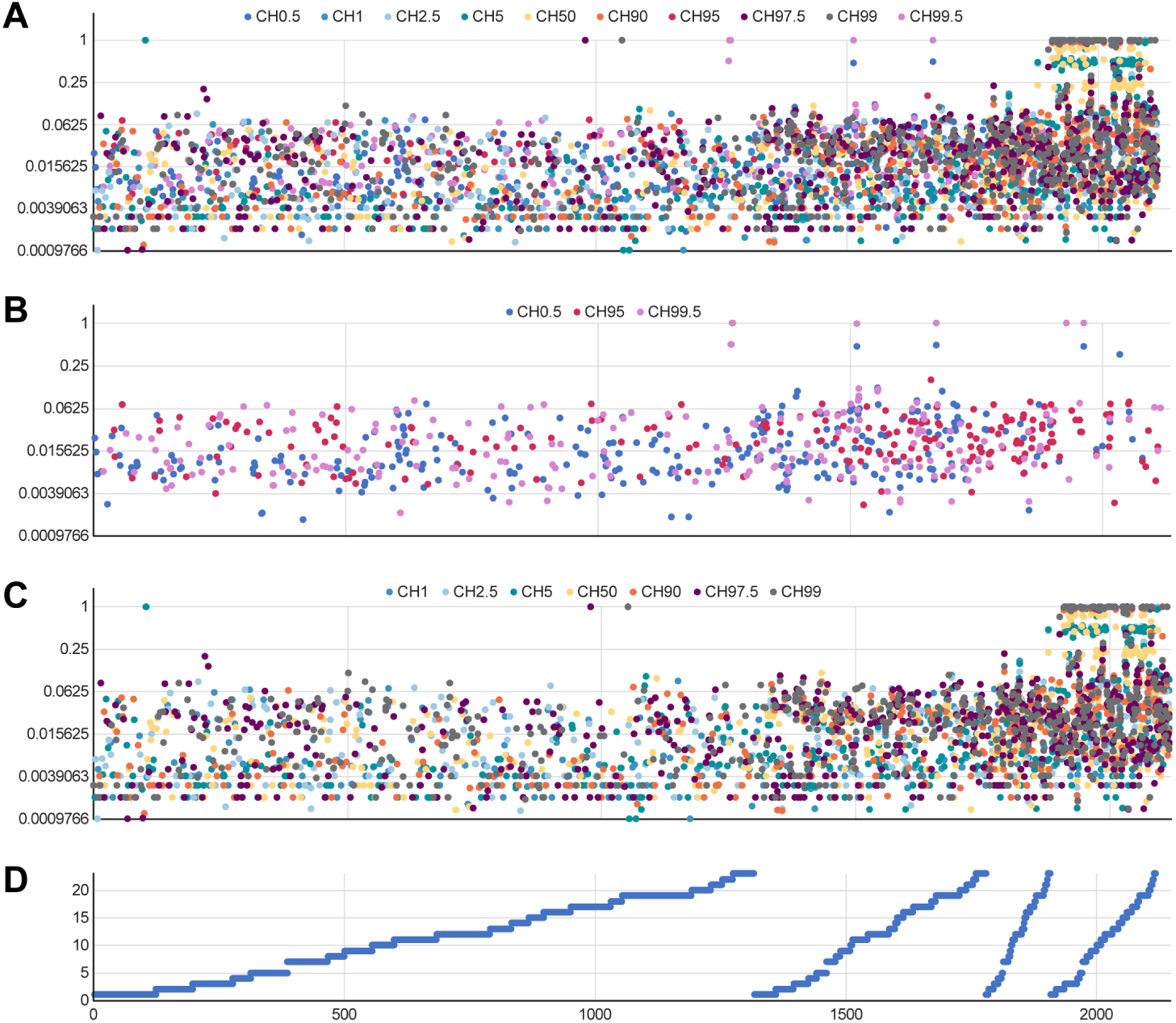
Positional distribution of FP NGS error alleles for the results from company CC by inhouse analysis. A. Total FP errors in DNA mixture samples. B. FP errors in 3 DNA mixture samples such as CH0.5, CH95, and CH99.5 which are the mixtures of DNA1 and DNA2 with the ratios of 99.5:0.5, 5:95, and 0.5:99.5, respectively. C. FP errors in the rest of DNA mixture samples except for CH0.5, CH95, and CH99.5. D. Chromosomal position of FP error alleles. Four portions depending on the frequency of errors at the specific allele sites were divided: 1st part from the left, alleles (N = 1316) with one FP error in 10 mixed reference standards; 2nd part, alleles with 2-3 FP errors (N = 462); 3rd part, allele with 4-5 FP errors (N = 119); 4th part, allele with 6 or more FP errors (N = 210). X-axis, chromosomal position. Y-axis, VAF of FP error variants.

### More transitional changes in FP errors with the employment of more sensitive condition (solid) for variant calls

Next, the transitional or transversional changes in the FP error alleles by default or solid mode of Dragen system were analyzed. With the default mode of Dragen system, relative rates of transitional or transversional changes were various depending on the companies: transition was higher in the results from companies AA and BB (Figs. 7A and 7C). In the results from company CC, changes of transition and transversion were similar (Fig. 7E). When the analyses were done by the solid mode of Dragen system which can detect more variants with lower VAF, the transition in FP error increased in companies AA, BB, and CC (Figs. 7B, 7D and 7F), suggesting that the efforts for sensitivity increase may produce more transition-related FP errors. The transitional changes by AF cutoff for company DD results were also observed (Figs. 7G and 7H), although not prominent as the solid condition analysis for the other companies AA, BB, and CC.

**Figure 7:**
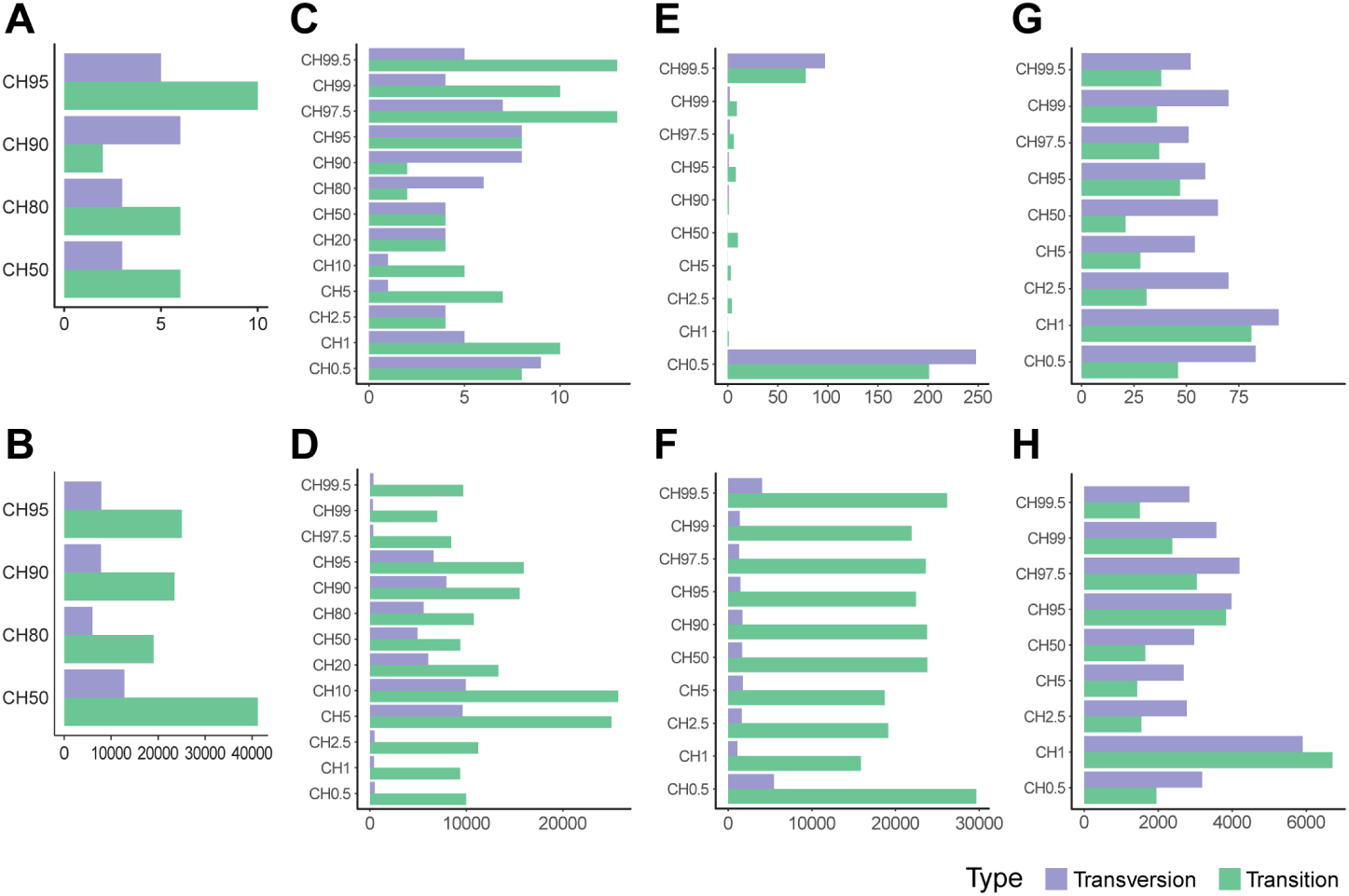
Transitional or transversional changes in FP errors. A. Variant analysis with default mode of Dragen system for the results from company AA. B. Variant analysis with solid mode of Dragen system for the results from company AA. C. Variant analysis with default mode of Dragen system for the results from company BB. D. Variant analysis with solid mode of Dragen system for the results from company BB. E. Variant analysis with default mode of Dragen system for the results from company CC. F. Variant analysis with solid mode of Dragen system for the results from company CC. G. Variant analysis after the selection of variants with AF *geq* 0.005 from the variant calls by DD inhouse method. H. Variant analysis with inhouse system of company DD. X-axis, the number of transition or transversion FP errors. Y-axis, reference-standard mixed DNAs. Sky blue, transversion. Green, transition.

### Little increase of sensitivity but big loss of specificity by the employment of more sensitive (solid) bioinformatic condition

The effects on the sensitivity and specificity by the analysis either default or solid Dragen condition for variant analyses were estimated. With the solid mode of Dragen system, 20 95 times more variants mostly with VAF less than 1% were called in DNA2 for companies AA, BB, and CC (Table S1). When the solid mode of Dragen system was employed for the bioinformatic analyses, the sensitivity increased 1.2 ∼1.8 times: 1.17 times in company AA, 1.84 times in company BB, and 1.57 times in company CC (Figs. 8A to 8C). However, the solid condition analyses increase FP errors dramatically (Fig. 8D): 8200 times in the result from company AA from 2.446 x 10^−6^ (median by default) to 2.005 x 10^−2^ (median by solid): 610 times in the result from company BB from 1.564 x 10^−5^ (median by default) to 9.543 x 10^−3^ (median by solid); 4152 times in the result from company CC from 1.230 x 10^−6^ (median by default) to 5.110 x 10^−3^ (median by solid). These results suggest that the adjustment of bioinformatic analysis conditions to increase sensitivity less than 2 times may lead to dramatic FP error increase (610∼8200 times).

**Figure 8:**
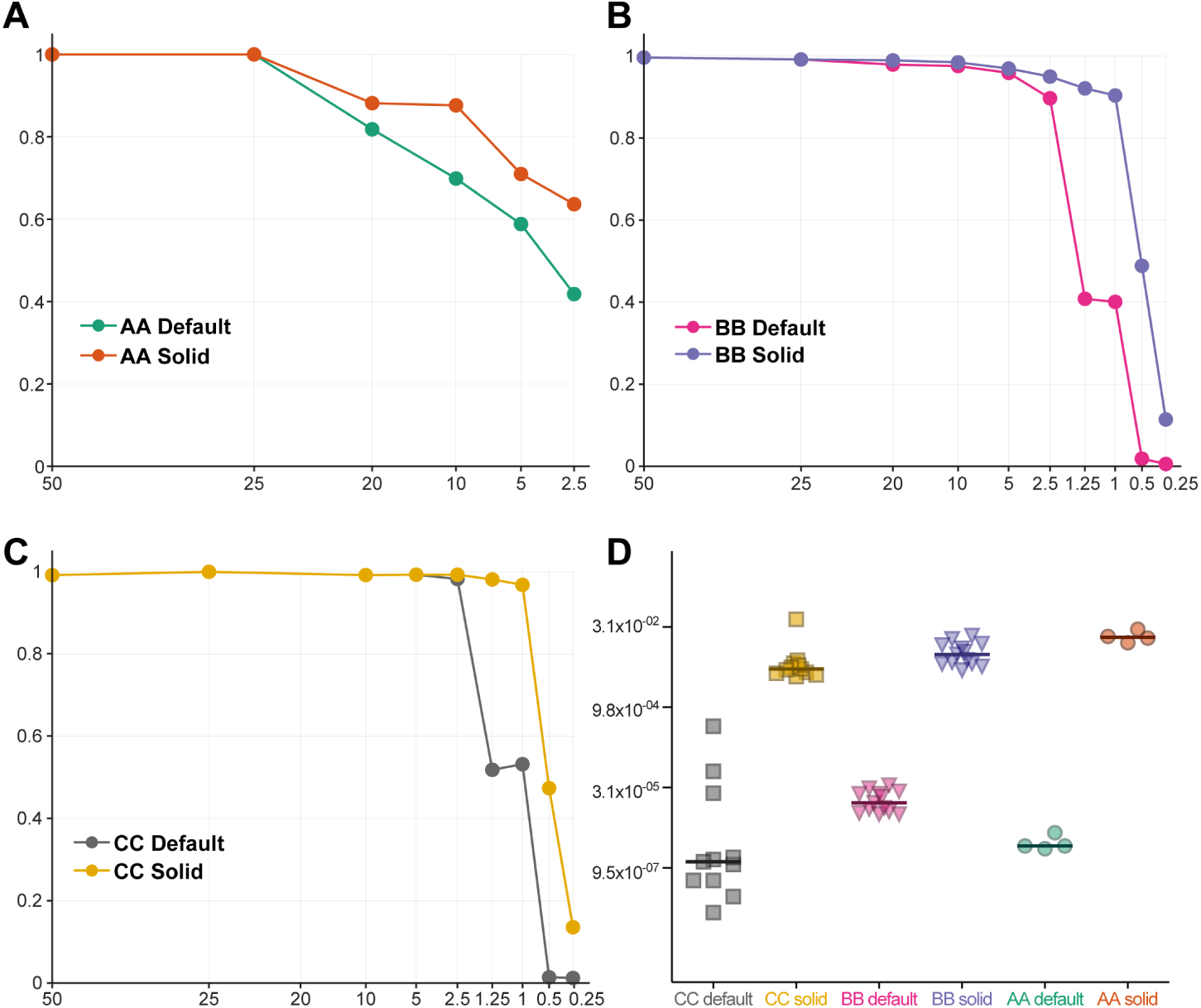
Difference in sensitivities and specificities depending on the analytic modes of Dragen system. Two modes such as default (less variants are called) and solid (more variants are called) modes were employed. A. Sensitivity difference in the results from company AA depending on the analytic modes (AA default, 24.66%; AA Solid, 21.01%). B. Sensitivity difference in the results from company BB depending on the analytic modes (BB default, 5.68%; BB Solid 3.08%). C. Sensitivity difference in the results from company CC depending on the analytic modes (CC default, 2.34%; CC Solid 1.49%). D. Specificity difference between two analytic modes in the results from AA, BB and CC companies. From figures A to C, X axis is the expected percentage of variant AF for informative alleles, and Y-axis is the detection rate of informative alleles at the expected percentage of VAF (X-axis). For D, Y-axis is FP error rate. For D X-axis, CC default, variants analyzed by default mode for the results from company CC; CC solid, variants analyzed by solid mode for the results from company CC; BB default, variants analyzed by default mode for the results from company BB; BB solid, variants analyzed by solid mode for the results from company BB; AA default, variants analyzed by default mode for the results from company AA; AA solid, variants analyzed by solid mode for the results from company AA.

## 3 Discussion

The sensitivity and specificity of NGS results from various service providers or kits have not been studied extensively [22]. Although NGS diagnosis, especially on cancer tissues, may have problems in the sensitivity and specificity, the possibility has been not fully estimated due to the limitations in evaluation methods [55, 39, 2, 58]. With our reference-standard materials consisting of various ratio mixtures of H. mole DNA and an individual’s blood genomic DNA, in the present study, the sensitivity and specificity for T-NGS results from certified service providers were evaluated: it was found that the sensitivity showed 16.3 times difference, and that the specificity showed about 4280 times difference.

Several reports have been published to estimate T-NGS sensitivity [12, 46, 9, 65, 7, 6], but more studies with newly developed methodologies are needed. In the present study, reference-standard materials consisting of various ratio mixtures of H. mole DNA and an individual’s blood genomic DNA were employed, and the sensitivity was evaluated by the analyses of the signals from diluted variants. In the analysis of NGS sensitivity, the variation was as large as 16.3 times from 1.51% to 24.7%. In the worst T-NGS results from company AA, 24.7% sensitivity of T-NGS suggests that only variants with over 25% allelic fraction can be reliably detected. Analysis with Dragen system for the results from company AA improved linearity of expected VAF versus VAF value from T-NGS results. However, the sensitivity from company AA was not improved by the analysis with Dragen system, suggesting that the raw data production was the main reason for the low sensitivity of the results from company AA, and the relatively low sequencing depth in the raw data may be one of main problems. However, other factors may also be involved, because the correlation between sequencing depth and sensitivity of T-NGS was not linear: the sensitivity of the results from company CC was about the same as that from company DD, although there was difference in sequencing depth. Also, the sensitivity of the results from company CC was better than that from company BB, although the sequencing depth was about the same. With the employment of Dragen system for bioinformatic analyses, the sensitivity was similar to that from inhouse methods (0.849 ∼1.140). These results suggest that the variation in T-NGS sensitivity is as large as 16.3 times but the reasons for the variation may be various in addition to those from the sequencing depth or the conditions for bioinformatic analyses.

The alleles in HLA region failed to show the dilution effect from the variant dilutions in the mixed DNA samples from company CC, but this kind of errors at specific alleles has not been reported, though the difficulty of bioinformatic analysis for HLA region has been reported [64, 61]. If the alleles in certain regions such as HLA region are in drug-response related genes, there might be a high risk of false negativity in reporting candidate markers. The reason why the dilution effects of HLA alleles were not observed in T-NGS is not clear, but the bioinformatic analyses of HLA region may not be as clear as the other regions [23, 28]. Therefore, employment of HLA region-specific bioinformatic methods such as HLAminer, HLAreporter, LATES, PHLAT, and Omixon Target HLA may be necessary.

Accurate variant calling in NGS data is a critical step [21, 31], and the detection of variants or mutations from individuals with genetic inheritance or cancer tissues was the main focus. Therefore, the calls for variants and mutations were very cautious and strict, but reference calls which are not critical for clinical decision may be permissive, leading to preferential reference base call. We found the biased preferential reference base call from the bioinformatic analysis in the present study: the FP errors were extremely high by the appearance of reference bases in V-V pairs. In addition, the analysis of the sensitivity with the employment of the diluted reference bases, instead of the diluted variants, showed unexpectedly high sensitivity: the diluted reference bases with less than 0.5% AF were found in almost all companies and methods. These results suggest the presence of biased preferential reference base call originated just from bioinformatic analyses. The significance of the preferential reference base calls from bioinformatic bias need more validation, but the biased calls should be considered as FP calls.

Few studies for the specificity of NGS have been reported [43, 39, 3, 52], but the routine estimation of FP error rates especially for low VAF alleles has not been performed due to the limitations of the evaluation methods. In the present study, the specificity of T-NGS was estimated with the employment of the reference standards. The variation in FP error rate was as large as 4280 times: the highest in the results for company DD (median: 5.814 x 10^−4^), and the lowest in those for company AA (median: 1.359 x 10^−7^). However, the lowest FP error rate of the results from company AA was probably related to extremely low sensitivity (only about 20% variant can be detected). The difference in FP error rate between company CC and DD was also as large as 473 (5.814 x 10^−4^/1.230 x 10^−6^). The big difference in the specificity of T-NGS is quite surprising because FP error may increase 4280 times or over 400 times depending on service providers and/or variant analysis methods. These results suggest that there are a lot of variations in the specificity even within the same Illumina platforms by certified NGS providers, which may hamper the right interpretation from T-NGS results for the selection of anti-cancer drugs based on precision medicine.

The T-NGS raw data generation for company DD adopted unique molecular identifier (UMI) to increase sensitivity and specificity [32, 60], whereas those for the other companies in the present study did not. In our results, the employment of UMI in the T-NGS raw data production by company DD showed better sensitivity than those by company AA and BB, but there was no sensitivity difference with company CC, suggesting that the NGS results without UMI may have sensitivity as high as UMI-employed NGS methods. Therefore, the benefit from the UMI employment in the NGS sensitivity should be carefully evaluated further, because questions on the benefit of UMI has been raised previously [4]. In our evaluation of specificity of T-NGS results for company DD, in addition, the FP error rate was much higher than those from company BB and CC which do not employ UMIs in their T-NGS raw data production, suggesting that the UMI employment may lead to high FP rates, in contrast to the general notion that the employment of UMIs can increase specificity [32, 63].

The NGS analysis was considered as a straightforward method, and some researchers published that NGS error is negligible [6, 68, 49]. Our previous study, however, showed that there are a lot of FN errors in whole exome or near whole exome NGS databases, which is confirmed by T-NGS analysis [30]. The present study results also indicate that there are a lot of FN and FP errors in T-NGS analyses, and also that there was an inverse correlation between sensitivity and specificity for T-NGS: the increase of the sensitivity as small as 2 times can lead to loss of the specificity as large as 59.5 times (as in company CC). In contrast, small sensitivity increase (less than 2 times) by the employment of more sensitive (solid) condition led to the big loss of specificity (FP error rates, 610 ∼8200 times) compared to less sensitive (default) one with the same Dragen system. In addition, the present study found batch to batch variations and sample to sample variations of specificity for T-NGS results which can be reduced by the employment of semiconductor NGS technology like Ion GeneStudio S5 System from Thermo Fisher. Furthermore, it was found that the adjustment of the analytic parameters with more sensitive conditions can lead to call 20 ∼95 times more variants, resulting in 8200 times more FP errors in the T-NGS results. These results suggest that the NGS analysis is not straightforward. Therefore, a lot of FP and FN errors in T-NGS results should be evaluated and the sincere efforts to reduce the errors both for the raw data generation and for the bioinformatic analyses should be pursued.

By the cooperative efforts of SEQC2 project, sensitivity and specificity of eight Pan-Cancer panels were evaluated [19] by the employment of genomic reference sample verified from another SEQC2 study [15]. In the Pan-Cancer panel evaluation study, the sensitivity which can be determined by the least percentage of allele fraction at which 95% of the diluted alleles, was about 1 ∼5%, and the best sensitivity results from the study is similar to those from the present study (1.51 ∼to 24.7%). Still, the variation of the sensitivity in various companies is larger in the present study, one of which reasons may be related to random selection of service providers, which performed the analysis along with the other samples, and which might have random errors or variations in their experimental process. The specificities of eight Pan-Cancer panels (10^−5^ ∼10^−6^ with the AF cutoff of 1%) from the SEQC2 project [19] seems to be similar to those from the present study (7.327 x 10^−5^ ∼1.230 x 10^−6^). However, the SEQC2 project [19] employed only about 50% of available informative alleles after pre-characterization of reference-standard materials by whole genome, whole exome sequencing, and digital droplet PCR methods, in contrast that the present study does not need those pre-characterization efforts on reference standards.

Researchers or service providers have already identified some of their own T-NGS-specific FP-error-prone sites during the analyses of NGS [48, 62, 10], but the presence of those error-prone sites for specific T-NGS kits has not been reported to the patients at the clinic or to the researchers. The present study showed the presence of FP-error prone sites, along with the identification method. The removal of error-prone alleles in the final reports can reduce FP errors significantly. On the other hand, the present study also showed the presence of the batch-to-batch error variations and the sample-to-sample error variations in a batch, suggesting the difficulty in removing the error-prone alleles. Our results indicate that more error-reducing efforts are needed before performing the T-NGS analyses, one of which may be the exclusion of FP errors in error-prone sites for which alleles should be reported to the researchers or clinicians at the same time.

The present study has limitations: 1) evaluation of the specificity and sensitivity only with the base changes, 2) using only freshly prepared DNA mixtures, 3) the employment of only Illumina systems for NGS analysis, and 4) due to the confidentiality, technical bioinformatic information from companies are not provided, which limits the detailed analyses of the reasons for the differences in sensitivity or specificity of the results from various companies. The present study analyzed only with base changes, but the sensitivity and specificity for the large segmental deletions or amplifications will be quite hard to evaluate, and reference-standard materials comprising diverse cells may be needed [17, 19]. The present study also has limitation in using only fresh DNAs: because clinical laboratories are using formalin-fixed paraffin-embedded (FFPE) methods to preserve solid tumor samples, and the sensitivity and specificity for the T-NGS results from FFPE tissues are needed. Because FFPE tissues have damaged DNA fragments and the modified bases which can confound the accurate variant calling even within a tumor tissue [69]. With the use of FFPE tissues or cells, the sensitivity and specificity of T-NGS would be lower than the ones from the present study with fresh DNA mixtures. For the applications in the clinics, FFPE cell block slides which can easily be prepared from the mixed cell lines with various ratios can be employed. In addition, many NGS platforms from various companies are currently employed, but few studies for the relative performance have been reported [16]: therefore, further validations on the various platforms for the analysis of alleles with low VAF are needed in the future.

The present study showed large variations in the sensitivity and specificity of T-NGS results even from the NGS certified service providers, and suggests that the quantitative measurements of the sensitivity and specificity may be essential for interpretation of T-NGS results at each laboratory and clinic. Recently, the difficulty in clinical implementation of the models from precision medicinal researches which prioritize novelty has been commented [38]. There should be a lot of reasons for the difficulties in the clinical implementation, but the complex procedure in the production and interpretation of NGS raw data and the lack of appropriate standardization protocols which are generally ignored or just have been recognized can be the most important reasons. Analysis of reference standards for NGS kits may be the big step toward the implementation of NGS results or kits to clinics. A large open source bioinformatic tools for variant calls as in Amazon Genomics CLI (https://aws.amazon.com/ko/genomics-cli/) or nf-core (https://nf-co.re/) are currently available [40], and the present study evaluation method for T-NGS may give a guide to choose better bioinformatic methods or conditions for the specific study purposes. For the quality evaluation of T-NGS kits and results at each laboratory, mixtures of homozygote DNA and blood DNA with various ratios can be easily employed as reference standards.

## 4 Methods

### Preparation of reference-standard DNA materials

An individual’s blood sample was obtained from with informed consent and this study was approved by the Institutional Review Board (IRB) of National Cancer Center, Korea. Genomic DNA from blood was extracted using DNeasy Blood and Tissue kit (Qiagen, Valencia, CA). Human H. mole cell DNA obtained from Coriell (NA07489, Camden, NJ) was purified again with QIAamp DNA mini kits to remove contaminated RNA. Mixtures of H. mole DNA (DNA1) and blood genomic DNA (DNA2), or reference-standard DNAs, were prepared with various ratios: CH0.5, mixture of blood DNA and H. mole with a ratio of 99.5:0.5; CH1, 99:1; CH2.5, 97.5:2.5; CH5, 95:5; CH10, 90:10; CH20, 80:20; CH50, 50:50; CH80, 20:80; CH90, 10:90; CH95, 5:95; CH97.5, 2.5:97.5; CH99, 1:99; CH99.5, 0.5:99.5. DNA concentration and purity was checked using a Nanodrop 8000 UV-vis spectrometer (Thermo Scientific, Waltham, MA) and Qubit 2.0 Fluorometer (Life Technologies, Grand Island, NY). The reference standards were sent to the companies AA, BB, and CC. The information of T-NGS kits for companies AA, BB, CC, and DD was shown in Table S2 and S3.

### Raw data production for T-NGS analyses

Quality and quantity of DNAs were checked again by each company’s own methods and criteria. Target capture, paired-end NGS sequencing library construction, and NGS sequencing were performed by each company’s own methods. All companies employed Illumina sequencing method for NGS sequencing. Companies AA and CC were accredited by CAP survey, and company BB by Korean FDA for T-NGS service. Raw data production employing the company DD’s kit which adopts unique molecular identifier (UMI), was performed by company CC.

### Variant detection and Alignment

Variants were analyzed by each company’s own bioinformatic method (inhouse method), and also by Dragen system (version 4.2, Illumina Inc., CA, USA). For the variant analyses by Dragen system, hg19 (https://www.ncbi.nlm.nih.gov/datasets/genome/GCF_000001405.13/) was employed as a reference genome. The bed file was provided only for company BB, and the variants only in exons of the target genes for companies AA, CC, and DD were analyzed. Either default or –vc-enable-umi-solid (solid) options for Dragen system was employed, and –vc-enable-umi-solid (solid) option was for the detection of more varaints with low allelic fraction. Annotation for Dragen system was performed by the employment of GATK Funcotator (FUNCtional annOTATOR). The inhouse analysis results from company DD were modified by selecting variant alleles only with VAF values *≥* 0.005, and the modified results from company DD were marked as DD AF *≥* 0.005.

The paired-end reads were aligned to the human reference genome (hg19) using BWA-MEM (v.0.7.5). SAMTOOLS (v0.1.18), GATK (v3.1-1), and Picard (v1.93) were used for file handling, local realignment, and removal of duplicate reads, respectively. We recalibrated base quality scores using GATK BaseRecalibrator based on known single-nucleotide polymorphisms (SNPs) and indels from dbSNP138.

### Estimation of T-NGS sensitivity

For NGS tests, detection sensitivity is typically specified in terms of Limit of Detection (LOD), following the guidelines established by The Clinical and Laboratory Standards Institute (ref, CLSI, 2012) or by American College of Medical Genetics and Genomics (ACMG) guideline for NGS [51], with the variant detection based on dilution assays. To estimate the sensitivity of T-NGS, Probit regression, a linear regression model with a cumulative normal distribution link function, was used to estimate a regression line similar to the graph using all or part of the observed values, and the LOD value was selected by estimating the lowest concentration that can detect 95% alleles.

For the evaluation of sensitivity, reference-standard DNAs, which are the mixtures of DNA1 (H. mole DNA) and DNA2 (blood DNA from an individual) at various ratios, were employed as reference standards for T-NGS. Among the variant calls from each company and/or method, informative alleles, in which variant or reference base is detected only one of DNA1 or DNA2, are selected. Three types of informative alleles were analyzed for sensitivity estimation: 1) N-Ho pair alleles which are null in DNA1 and homozygote variant in DNA2; 2) Ho-N pair alleles which are homozygote variant in DNA1 and null in DNA2; and 3) N-He pair alleles which are null in DNA1 and heterozygote in DNA2. In the selection of the informative allele types in DNA1 or DNA2, variant alleles with VAF < 0.05 were considered as null; alleles with VAF > 0.95 as homozygote; and alleles with 0.3 < VAF < 0.7 as heterozygote. After the determination of null, homozygote, or variant alleles in DNA1 and DNA2, the informative allele types such as N-Ho, Ho-N, and N-He pairs were determined. Then, the absence of the variant in the DNA mixtures at the informative alleles were considered as false negative, and the VAF values of informative alleles present in the DNA mixtures were employed for the evaluation of the linearity with the dilution fold of the variants.

### Estimation of T-NGS specificity

For the evaluation of specificity, reference-standard DNAs were also employed. FP error alleles were defined as the alleles with the base(s) from the T-NGS results in mixed reference-standard DNAs in the absence of that base(s) from the T-NGS results in either DNA1 or DNA2. FP errors occurs in three pairs such as 1) R-R pair alleles at which DNA1 and DNA2 have reference base; 2) V-V pair alleles at which neither DNA1 nor DNA2 have reference base; and 3) V-R pair alleles at which DNA1 and DNA2 have both reference and variant bases. FP errors are identified in T-NGS results from the mixed DNA samples. In DNA1 and DNA2 for R-R pair alleles, the variant alleles are absent. The alleles are considered as V-V or V-R pair if any variant or reference is present in DNA1 or DNA2 for V-V or V-R pair alleles. The appearance of reference or variant bases which are not present in DNA1 or DNA2, was considered as FP error. The FP errors by the appearance of reference bases in the DNA mixtures at the V-V pair alleles were defined as VVR errors which were removed for the calculation of total FP errors. The FP error rates were calculated as the FP errors divided by total pair allele number in exons of target genes for each T-NGS kit.

## Availability of data and materials

All T-NGS sequencing fastq files used for this study are publicly available in the NCBI Sequence Read Archive (SRA) under the BioProject accession number PRJNA1134909.

## Supplementary data

Supplementary data to this article can be found in the Supplementary date file.

## Acknowledgements

The service for bioinformatic analyses with Dragen system from raw data was provided by Bioinformatics Analysis Team, and statistical analyses were performed by Biostatistics Collaboration Team of Research Core Center at National Cancer Center, Korea.

## Funding

This research was supported by the intramural program of National Cancer Center, Korea (2210700 and 2311430 to K-M. H).

## Notes

### Competing Interest Statement

The authors have declared no competing interest.

## References

[1] A. Abbasi and L. B. Alexandrov. Significance and limitations of the use of next-generation sequencing technologies for detecting mutational signatures. DNA Repair (Amst), 107:103200, 2021.

[2] U. Bacher, E. Shumilov, J. Flach, N. Porret, R. Joncourt, G. Wiedemann, M. Fiedler, U. Novak, U. Amstutz, and T. Pabst. Challenges in the introduction of next-generation sequencing (ngs) for diagnostics of myeloid malignancies into clinical routine use. Blood Cancer J, 8:113, 2018.

[3] P. Bauer, K. K. Kandaswamy, M. E. R. Weiss, O. Paknia, M. Werber, A. M. Bertoli-Avella, Z. Yuksel, M. Bochinska, G. E. Oprea, S. Kishore, et al. Development of an evidence-based algorithm that optimizes sensitivity and specificity in es-based diagnostics of a clinically heterogeneous patient population. Genet Med, 21:53–61, 2019.

[4] J. Bieler, S. Kubik, M. Macheret, C. Pozzorini, A. Willig, and Z. Xu. Benefits of applying molecular barcoding systems are not uniform across different genomic applications. J Transl Med, 21:305, 2023.

[5] W. Chen, Y. Zhao, X. Chen, Z. Yang, X. Xu, Y. Bi, V. Chen, J. Li, H. Choi, B. Ernest, et al. A multicenter study benchmarking single-cell rna sequencing technologies using reference samples. Nat Biotechnol, 39:1103–1114, 2021.

[6] C. Cheng, Z. Fei, and P. Xiao. Methods to improve the accuracy of next-generation sequencing. Front Bioeng Biotechnol, 11:982111, 2023.

[7] C. Cheng and P. Xiao. Evaluation of the correctable decoding sequencing as a new powerful strategy for dna sequencing. Life Sci Alliance, 5, 2022.

[8] ITP-CAoWG Consortium. Pan-cancer analysis of whole genomes. Nature, 578:82–93, 2020.

[9] P. Dai, L. R. Wu, S. X. Chen, M. X. Wang, L. Y. Cheng, J. X. Zhang, P. Hao, W. Yao, J. Zarka, G. C. Issa, et al. Calibration-free ngs quantitation of mutations below 0.01 Nat Commun, 12:6123, 2021.

[10] E. M. Davis, Y. Sun, Y. Liu, P. Kolekar, Y. Shao, K. Szlachta, H. L. Mulder, D. Ren, S. V. Rice, Z. Wang, et al. Sequencerr: measuring and suppressing sequencer errors in next-generation sequencing data. Genome Biol, 22:37, 2021.

[11] D. Deshpande, K. Chhugani, Y. Chang, A. Karlsberg, C. Loeffler, J. Zhang, A. Muszynska, V. Munteanu, H. Yang, J. Rotman, et al. Rna-seq data science: From raw data to effective interpretation. Front Genet, 14:997383, 2023.

[12] I. W. Deveson, B. Gong, K. Lai, J. S. LoCoco, T. A. Richmond, J. Schageman, Z. Zhang, N. Novoradovskaya, J. C. Willey, W. Jones, et al. Evaluating the analytical validity of circulating tumor dna sequencing assays for precision oncology. Nat Biotechnol, 39:1115–1128, 2021.

[13] L. T. Fang, B. Zhu, Y. Zhao, W. Chen, Z. Yang, L. Kerrigan, K. Langenbach, M. de Mars, C. Lu, K. Idler, et al. Establishing community reference samples, data and call sets for benchmarking cancer mutation detection using whole-genome sequencing. Nat Biotechnol, 39:1151–1160, 2021.

[14] R. Fisher, L. Pusztai, and C. Swanton. Cancer heterogeneity: implications for targeted therapeutics. Br J Cancer, 108:479–485, 2013.

[15] U.S. Food and Drug Administration-(FDA). Considerations for design, development, and analytical validation of next generation sequencing (ngs) – based in vitro diagnostics (ivds) intended to aid in the diagnosis of suspected germline diseases, 2020. https://www.fda.gov/regulatory-information/search-fda-guidance-documents/considerations-design-development-and-analytical-validation-next-generation-sequencing-ngs-based.

[16] J. Foox, S. W. Tighe, C. M. Nicolet, J. M. Zook, M. Byrska-Bishop, W. E. Clarke, M. M. Khayat, M. Mahmoud, P. K. Laaguiby, Z. T. Herbert, et al. Performance assessment of dna sequencing platforms in the abrf next-generation sequencing study. Nat Biotechnol, 39:1129–1140, 2021.

[17] G. M. Frampton, A. Fichtenholtz, G. A. Otto, K. Wang, S. R. Downing, J. He, M. Schnall-Levin, J. White, E. M. Sanford, P. An, et al. Development and validation of a clinical cancer genomic profiling test based on massively parallel dna sequencing. Nat Biotechnol, 31:1023–1031, 2013.

[18] B. Gong, R. Kusko, W. Jones, W. Tong, and J. Xu. Ultra-deep multi-oncopanel sequencing of benchmarking samples with a wide range of variant allele frequencies. Sci Data, 9:288, 2022.

[19] B. Gong, D. Li, R. Kusko, N. Novoradovskaya, Y. Zhang, S. Wang, C. Pabon-Pena, Z. Zhang, K. Lai, W. Cai, et al. Cross-oncopanel study reveals high sensitivity and accuracy with overall analytical performance depending on genomic regions. Genome Biol, 22:109, 2021.

[20] S. Goodwin, J. D. McPherson, and W. R. McCombie. Coming of age: ten years of next-generation sequencing technologies. Nat Rev Genet, 17:333–351, 2016.

[21] C. Groza, T. Kwan, N. Soranzo, T. Pastinen, and G. Bourque. Personalized and graph genomes reveal missing signal in epigenomic data. Genome Biol, 21:124, 2020.

[22] O. Harismendy, P. C. Ng, R. L. Strausberg, X. Wang, T. B. Stockwell, K. Y. Beeson, N. J. Schork, S. S. Murray, E. J. Topol, S. Levy, and K. A. Frazer. Evaluation of next generation sequencing platforms for population targeted sequencing studies. Genome Biol, 10:R32, 2009.

[23] K. Hosomichi, T. Shiina, A. Tajima, and I. Inoue. The impact of next-generation sequencing technologies on hla research. J Hum Genet, 60:665–673, 2015.

[24] M. Jang, H. Y. Pak, J. Y. Heo, H. Lim, Y. L. Choi, H. S. Shim, and E. K. Kim. Trends and clinical characteristics of next-generation sequencing-based genetic panel tests: An analysis of korean nationwide claims data. Cancer Res Treat, 56:27–36, 2024.

[25] L. J. Jennings, M. E. Arcila, C. Corless, S. Kamel-Reid, I. M. Lubin, J. Pfeifer, R. L. Temple-Smolkin, K. V. Voelkerding, and M. N. Nikiforova. Guidelines for validation of next-generation sequencing-based oncology panels: A joint consensus recommendation of the association for molecular pathology and college of american pathologists. J Mol Diagn, 19:341–365, 2017.

[26] W. Jones, B. Gong, N. Novoradovskaya, D. Li, R. Kusko, T. A. Richmond, D. J. Jr. Johann, H. Bisgin, S. M. E. Sahraeian, P. R. Bushel, et al. A verified genomic reference sample for assessing performance of cancer panels detecting small variants of low allele frequency. Genome Biol, 22:111, 2021.

[27] K. Jung, S. Lee, H. Y. Na, J. W. Kim, J. C. Lee, J. H. Hwang, J. W. Kim, and J. Kim. Ngs-based targeted gene mutational profiles in korean patients with pancreatic cancer. Sci Rep, 12:20937, 2022.

[28] S. Ka, S. Lee, J. Hong, Y. Cho, J. Sung, H. N. Kim, H. L. Kim, and J. Jung. Hlascan: genotyping of the hla region using next-generation sequencing data. BMC Bioinformatics, 18:258, 2017.

[29] H. Kim, J. W. Yun, S. T. Lee, H. J. Kim, S. H. Kim, J. W. Kim, and Korean Society for Genetic Diagnostics Clinical Guidelines C. Korean society for genetic diagnostics guidelines for validation of next-generation sequencing-based somatic variant detection in hematologic malignancies. Ann Lab Med, 39:515–523, 2019.

[30] Y. H. Kim, Y. Song, J. K. Kim, T. M. Kim, H. W. Sim, H. L. Kim, H. Jang, Y. W. Kim, and K. M. Hong. False-negative errors in next-generation sequencing contribute substantially to inconsistency of mutation databases. PLoS One, 14:e0222535, 2019.

[31] D. C. Koboldt. Best practices for variant calling in clinical sequencing. Genome Med, 12:91, 2020.

[32] R. Kou, H. Lam, H. Duan, L. Ye, N. Jongkam, W. Chen, S. Zhang, and S. Li. Benefits and challenges with applying unique molecular identifiers in next generation sequencing to detect low frequency mutations. PLoS One, 11:e0146638, 2016.

[33] J. Lee, S. Choi, D. Jung, Y. Jung, J. H. Kim, S. Jung, and W. S. Lee. Mutational characterization of colorectal cancer from korean patients with targeted sequencing. J Cancer, 12:7300–7310, 2021.

[34] S. H. Lee, B. Lee, J. H. Shim, K. W. Lee, J. W. Yun, S. Y. Kim, T. Y. Kim, Y. H. Kim, Y. H. Ko, H. C. Chung, et al. Landscape of actionable genetic alterations profiled from 1,071 tumor samples in korean cancer patients. Cancer Res Treat, 51:211–222, 2019.

[35] H. Li. Improving snp discovery by base alignment quality. Bioinformatics, 27:1157–1158, 2011.

[36] W. Li, X. Huang, R. Patel, E. Schleifman, S. Fu, D. S. Shames, and J. Zhang. Analytical evaluation of circulating tumor dna sequencing assays. Sci Rep, 14:4973, 2024.

[37] F. Luh and Y. Yen. Fda guidance for next generation sequencing-based testing: balancing regulation and innovation in precision medicine. NPJ Genom Med, 3:28, 2018.

[38] F. Markowetz. All models are wrong and yours are useless: making clinical prediction models impactful for patients. NPJ Precis Oncol, 8:54, 2024.

[39] G. Matthijs, E. Souche, M. Alders, A. Corveleyn, S. Eck, I. Feenstra, V. Race, E. Sistermans, M. Sturm, M. Weiss, et al. Guidelines for diagnostic next-generation sequencing. Eur J Hum Genet, 24:2–5, 2016.

[40] T. R. Mercer, J. Xu, C. E. Mason, W. Tong, and M. S. Consortium. The sequencing quality control 2 study: establishing community standards for sequencing in precision medicine. Genome Biol, 22:306, 2021.

[41] T. Miura, S. Yasuda, and Y. Sato. A simple method to estimate the in-house limit of detection for genetic mutations with low allele frequencies in whole-exome sequencing analysis by next-generation sequencing. BMC Genom Data, 22:8, 2021.

[42] L. Moore, A. Cagan, T. H. H. Coorens, M. D. C. Neville, R. Sanghvi, M. A. Sanders, T. R. W. Oliver, D. Leongamornlert, P. Ellis, A. Noorani, et al. The mutational landscape of human somatic and germline cells. Nature, 597:381–386, 2021.

[43] W. Mu, H. M. Lu, J. Chen, S. Li, and A. M. Elliott. Sanger confirmation is required to achieve optimal sensitivity and specificity in next-generation sequencing panel testing. J Mol Diagn, 18:923–932, 2016.

[44] J. Nangalia and P. J. Campbell. Genome sequencing during a patient’s journey through cancer. N Engl J Med, 381:2145–2156, 2019.

[45] K. N. Natarajan, Z. Miao, M. Jiang, X. Huang, H. Zhou, J. Xie, C. Wang, S. Qin, Z. Zhao, L. Wu, et al. Comparative analysis of sequencing technologies for single-cell transcriptomics. Genome Biol, 20:70, 2019.

[46] A. M. Newman, S. V. Bratman, J. To, J. F. Wynne, N. C. Eclov, L. A. Modlin, C. L. Liu, J. W. Neal, H. A. Wakelee, R. E. Merritt, et al. An ultrasensitive method for quantitating circulating tumor dna with broad patient coverage. Nat Med, 20:548–554, 2014.

[47] A. Petrackova, M. Vasinek, L. Sedlarikova, T. Dyskova, P. Schneiderova, T. Novosad, T. Papajik, and E. Kriegova. Standardization of sequencing coverage depth in ngs: Recommendation for detection of clonal and subclonal mutations in cancer diagnostics. Front Oncol, 9:851, 2019.

[48] S. P. Pfeifer. From next-generation resequencing reads to a high-quality variant data set. Heredity (Edinb), 118:111–124, 2017.

[49] F. Pfeiffer, C. Grober, M. Blank, K. Handler, M. Beyer, J. L. Schultze, and G. Mayer. Systematic evaluation of error rates and causes in short samples in next-generation sequencing. Sci Rep, 8:10950, 2018.

[50] M. A. Quail, M. Smith, P. Coupland, T. D. Otto, S. R. Harris, T. R. Connor, A. Bertoni, H. P. Swerdlow, and Y. Gu. A tale of three next generation sequencing platforms: comparison of ion torrent, pacific biosciences and illumina miseq sequencers. BMC Genomics, 13:341, 2012.

[51] H. L. Rehm, S. J. Bale, P. Bayrak-Toydemir, J. S. Berg, K. K. Brown, J. L. Deignan, M. J. Friez, B. H. Funke, M. R. Hegde, E. Lyon, et al. Acmg clinical laboratory standards for next-generation sequencing. Genet Med, 15:733–747, 2013.

[52] A. Rose Brannon, G. Jayakumaran, M. Diosdado, J. Patel, A. Razumova, Y. Hu, F. Meng, M. Haque, J. Sadowska, B. J. Murphy, et al. Enhanced specificity of clinical high-sensitivity tumor mutation profiling in cell-free dna via paired normal sequencing using msk-access. Nat Commun, 12:3770, 2021.

[53] A. Rubben and A. Araujo. Cancer heterogeneity: converting a limitation into a source of biologic information. J Transl Med, 15:190, 2017.

[54] S. M. E. Sahraeian, L. T. Fang, K. Karagiannis, M. Moos, S. Smith, L. Santana-Quintero, C. Xiao, M. Colgan, H. Hong, M. Mohiyuddin, and W. Xiao. Achieving robust somatic mutation detection with deep learning models derived from reference data sets of a cancer sample. Genome Biol, 23:12, 2022.

[55] J. J. Salk, M. W. Schmitt, and L. A. Loeb. Enhancing the accuracy of next-generation sequencing for detecting rare and subclonal mutations. Nat Rev Genet, 19:269–285, 2018.

[56] H. T. Shin, Y. L. Choi, J. W. Yun, N. K. D. Kim, S. Y. Kim, H. J. Jeon, J. Y. Nam, C. Lee, D. Ryu, S. C. Kim, et al. Prevalence and detection of low-allele-fraction variants in clinical cancer samples. Nat Commun, 8:1377, 2017.

[57] B. E. Slatko, A. F. Gardner, and F. M. Ausubel. Overview of next-generation sequencing technologies. Curr Protoc Mol Biol, 122:e59, 2018.

[58] P. Song, L. R. Wu, Y. H. Yan, J. X. Zhang, T. Chu, L. N. Kwong, A. A. Patel, and D. Y. Zhang. Limitations and opportunities of technologies for the analysis of cell-free dna in cancer diagnostics. Nat Biomed Eng, 6:232–245, 2022.

[59] K. J. Suh, S. H. Kim, Y. J. Kim, H. Shin, E. Kang, E. K. Kim, S. Lee, J. W. Woo, H. Y. Na, S. Ahn, et al. Clinical application of next-generation sequencing in patients with breast cancer: Real-world data. J Breast Cancer, 25:366–378, 2022.

[60] J. Sun, M. Philpott, D. Loi, S. Li, P. Monteagudo-Mesas, G. Hoffman, J. Robson, N. Mehta, V. Gamble, T. Jr. Brown, et al. Correcting pcr amplification errors in unique molecular identifiers to generate accurate numbers of sequencing molecules. Nat Methods, 21:401–405, 2024.

[61] A. Sverchkova, S. Burkholz, R. Rubsamen, R. Stratford, and T. Clancy. Integrative hla typing of tumor and adjacent normal tissue can reveal insights into the tumor immune response. BMC Med Genomics, 17:37, 2024.

[62] M. Tomkova and B. Schuster-Bockler. Dna modifications: Naturally more error prone? Trends Genet, 34:627–638, 2018.

[63] M. Tsagiopoulou, M. C. Maniou, N. Pechlivanis, A. Togkousidis, M. Kotrova, T. Hutzenlaub, I. Kappas, A. Chatzidimitriou, and F. Psomopoulos. Umic: A preprocessing method for umi deduplication and reads correction. Front Genet, 12:660366, 2021.

[64] E. T. Weimer, M. Montgomery, R. Petraroia, J. Crawford, and J. L. Schmitz. Performance characteristics and validation of next-generation sequencing for human leucocyte antigen typing. J Mol Diagn, 18:668–675, 2016.

[65] J. C. Willey, T. B. Morrison, B. Austermiller, E. L. Crawford, D. J. Craig, T. M. Blomquist, W. D. Jones, A. Wali, J. S. Lococo, N. Haseley, et al. Advancing ngs quality control to enable measurement of actionable mutations in circulating tumor dna. Cell Rep Methods, 1:100106, 2021.

[66] W. Xiao, L. Ren, Z. Chen, L. T. Fang, Y. Zhao, J. Lack, M. Guan, B. Zhu, E. Jaeger, L. Kerrigan, et al. Toward best practice in cancer mutation detection with whole-genome and whole-exome sequencing. Nat Biotechnol, 39:1141–1150, 2021.

[67] J. Zhang, S. S. Spath, S. L. Marjani, W. Zhang, and X. Pan. Characterization of cancer genomic heterogeneity by next-generation sequencing advances precision medicine in cancer treatment. Precis Clin Med, 1:29–48, 2018.

[68] T. H. Zhang, N. C. Wu, and R. Sun. A benchmark study on error-correction by read-pairing and tag-clustering in amplicon-based deep sequencing. BMC Genomics, 17:108, 2016.

[69] Y. Zhang, T. M. Blomquist, R. Kusko, D. Stetson, Z. Zhang, L. Yin, R. Sebra, B. Gong, J. S. Lococo, V. K. Mittal, et al. Deep oncopanel sequencing reveals within block position-dependent quality degradation in ffpe processed samples. Genome Biol, 23:141, 2022.

[70] Y. Zhao, L. T. Fang, T. W. Shen, S. Choudhari, K. Talsania, X. Chen, J. Shetty, Y. Kriga, B. Tran, B. Zhu, et al. Whole genome and exome sequencing reference datasets from a multi-center and cross-platform benchmark study. Sci Data, 8:296, 2021.

