## Supplementary file for "Evaluation of false positive and false negative errors in targeted next generation sequencing"

Comparison of false positive and false negative errors in targeted next generation sequencing results from certified NGS service providers

Youngbeen Moon ^1†^, Young-Ho Kim^2†^, Jong-Kwang Kim^1^, Chung Hwan Hong^3^, Eun-Kyung Kang^3^, Hye Won Choi^3^, Dong‑eun Lee^4^, Tae-Min Kim^5^, Seong Gu Heo^6,7,8^, Namshik Han ^9,10,11^, Kyeong-Man Hong^1,3*^

^1^Bioinformatics Analysis Team, Research Core Center, Research Institute, National Cancer Center, Goyang, Gyeonggi‑do, Republic of Korea.

^2^Diagnostic and Therapeutics Technology Branch, Division of Technology Convergence, Research Institute, National Cancer Center, Goyang, Gyeonggi‑do, Republic of Korea.

^3^Cancer Molecular Biology Branch, Division of Cancer Biology, Research Institute, National Cancer Center, Goyang, Gyeonggi‑do, Republic of Korea.

^4^Biostatistics Collaboration Team, Research Core Center, Research Institute, National Cancer Center, Goyang, Gyeonggi‑do, Republic of Korea.

^5^Department of Medical Informatics and Cancer Research Institute, College of Medicine, The Catholic University of Korea, Seoul, Korea.

^6^Dana Farber Cancer Institute, Boston, MA, USA

^7^The Broad Institute of MIT and Harvard, Cambridge, MA, USA

^8^Harvard Medical School, Boston, MA, USA.

^9^Milner Therapeutics Institute, University of Cambridge, Cambridge, UK

^10^Cambridge Centre for AI in Medicine, Department of Applied Mathematics and Theoretical Physics, University of Cambridge, Cambridge, UK

^11^Wellcome Trust-Medical Research Council Cambridge Stem Cell Institute, University of Cambridge, Cambridge, UK.

**Table of contents**

**Supplementary figures S1–S7 and their legends**

**Supplementary table1-3**

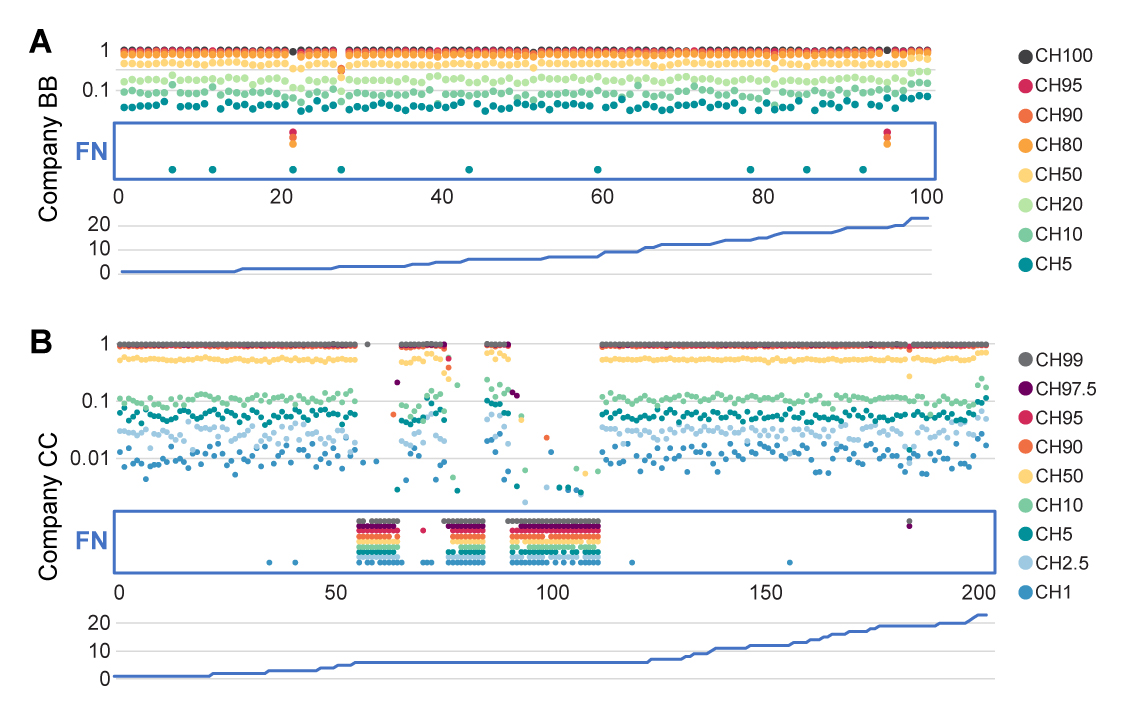
**Fig. S1** FN errors of T-NGS results from companies BB and CC in the alleles with Ho-N pair alleles. A. FN errors in T-NGS results from company BB. B. FN errors in T-NGS results from company CC. A lot of Ho-N pair alleles were FN at the alleles in chromosome 6. FN alleles at mixed reference standards were marked by red rectangle. At the lower columns, chromosome numbers were shown. The ratios of DNA1 and DNA2 for each sample (CH0 ~ CH100) were shown in Materials and Methods. X-axis, informative alleles aligned by chromosome and position. Y-axis, variant allelic fraction of informative alleles.

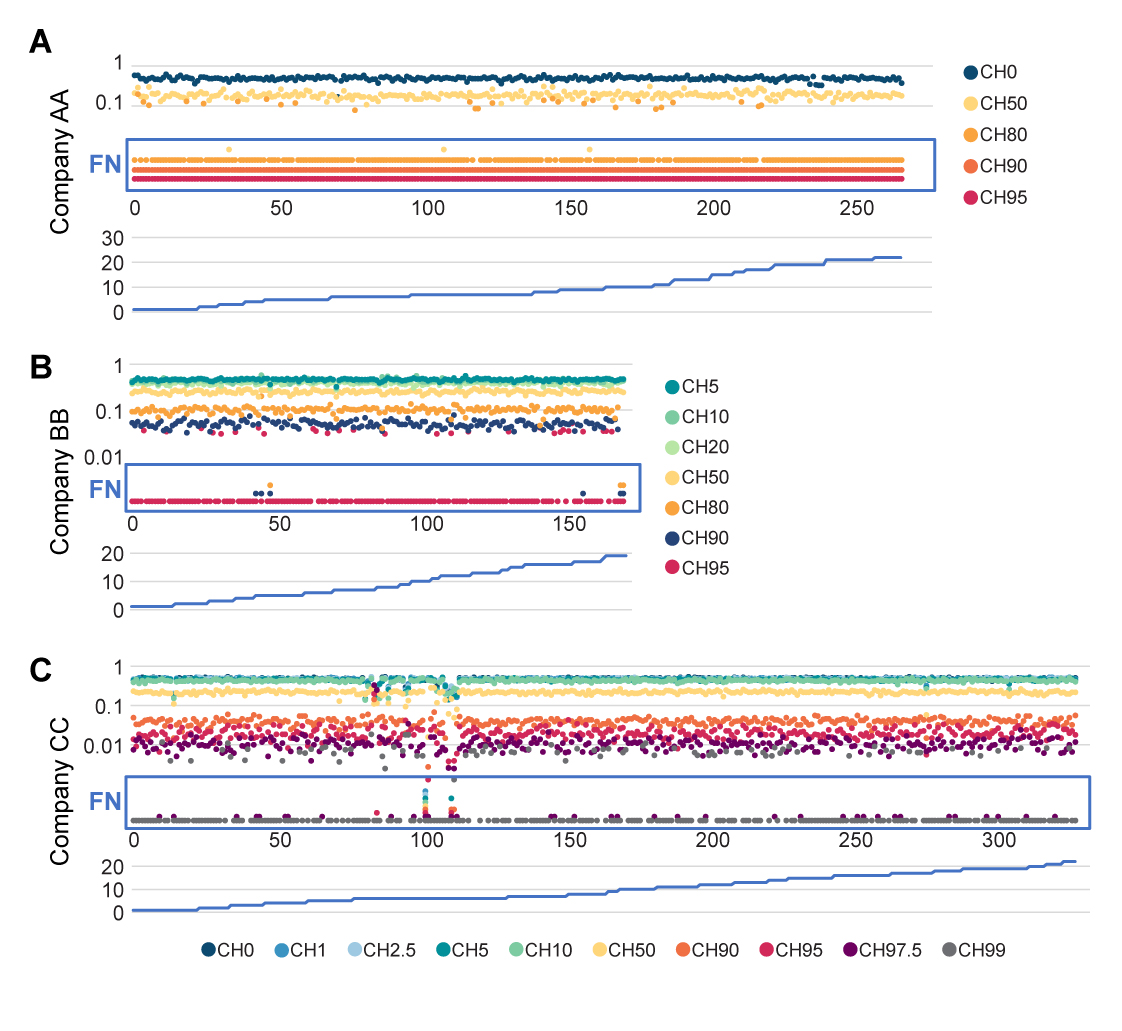

**Fig. S2** FN errors of T-NGS results from companies BB and CC in the He-N pair alleles. A. FN errors in T-NGS results from company AA. B. FN errors in T-NGS results from company BB. C. FN errors in T-NGS results from company CC. The informative alleles in HLA region of chromosome 6 were false negative. False negative alleles at mixed reference standards were shown and marked in FN column (red rectangle). At the lower columns, chromosome numbers were shown. The ratios of DNA1 and DNA2 for each sample (CH0 ~ CH100) were shown in Materials and Methods. X-axis, informative alleles aligned by chromosome and position. Y-axis, variant allelic fraction of informative alleles.

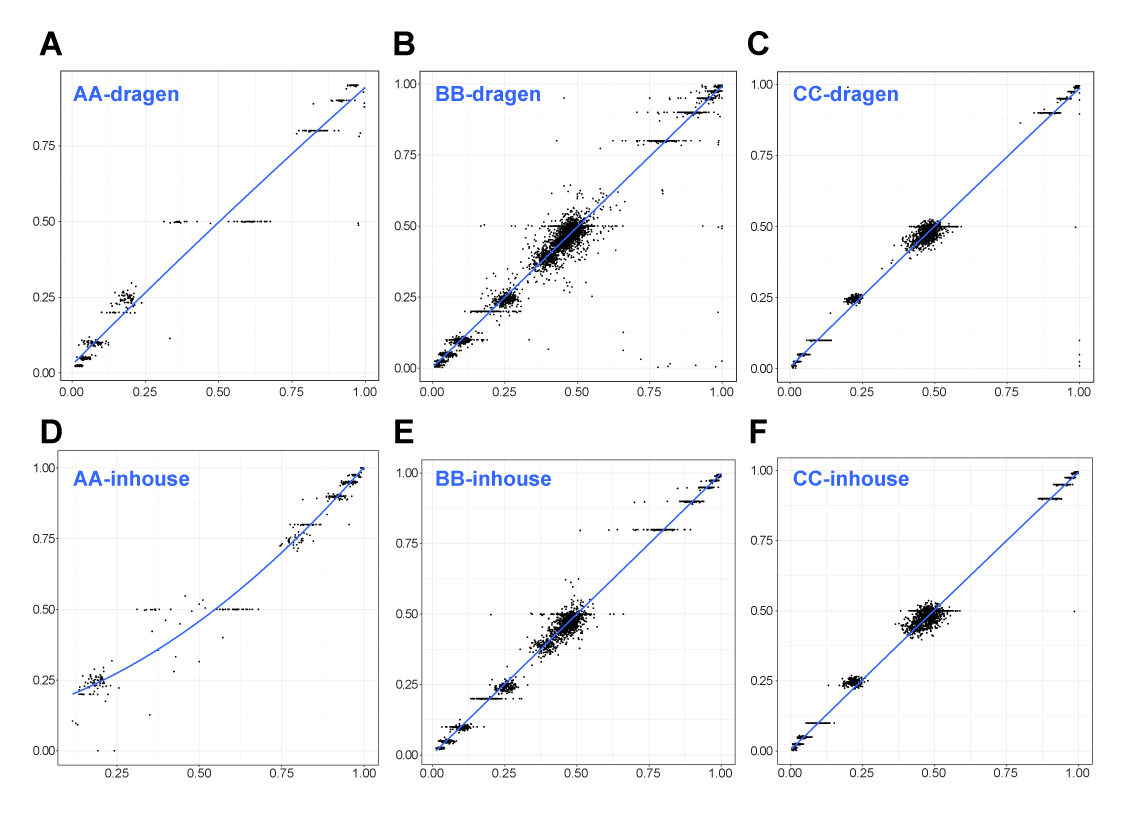

**Fig. S3** Correlation of allelic fraction values from T-NGS analysis with the expected allelic fraction values depending on the analysis methods. A. The correlation of allelic fractions for variants analyzed by Dragen system on the fastq files from company AA (AA-dragen, R = 0.9856). B. The correlation of allelic fractions for variants analyzed by Dragen system on the fastq files from company BB (BB-dragen, R = 0.9924). C. The correlation of allelic fractions for variants analyzed by Dragen system on the fastq files from CC (CC-dragen, R = 0.9957). D. The correlation of allelic fractions for variants analyzed by company AA’s inhouse method (AA-inhouse, R = 0.9789). E. The correlation of allelic fractions for variants analyzed by company BB’s inhouse method (BB-inhouse, R = 0.9969). F. The correlation of allelic fractions for variants analyzed by company CC’s inhouse method (CC-inhouse, R = 0.9935). X-axis, allelic fraction values from T-NGS analysis. Y-axis, the expected allelic fraction value.

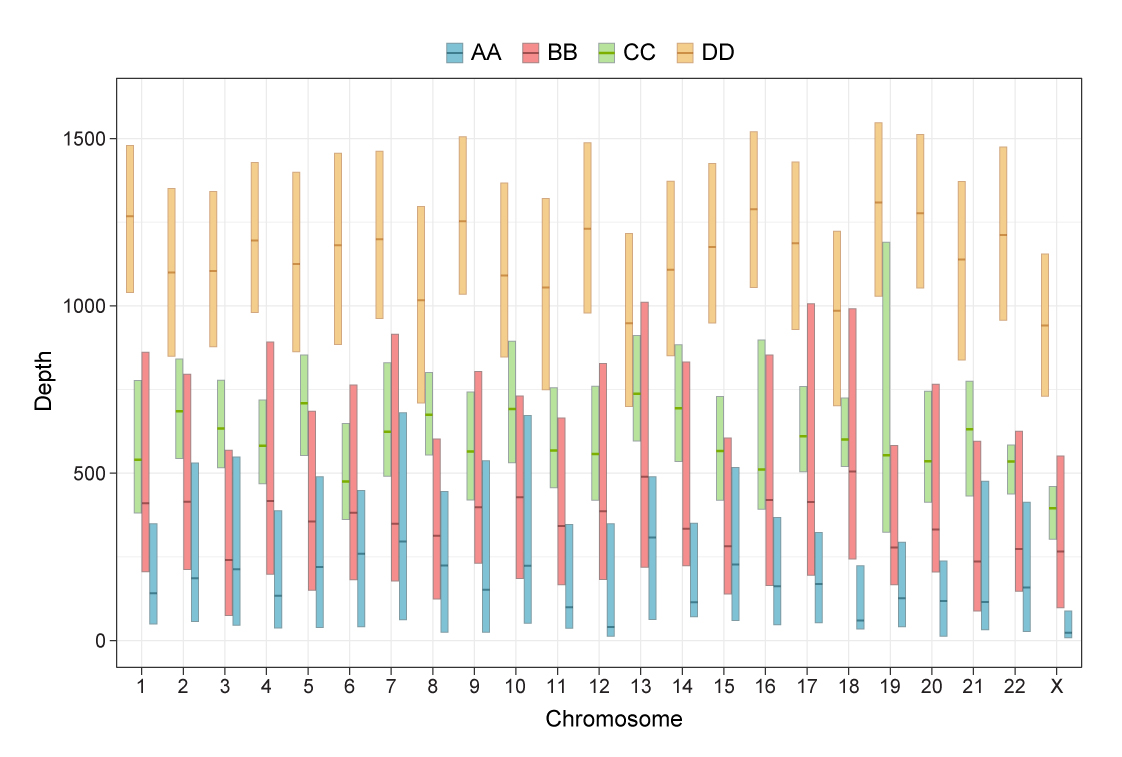

**Fig. S4** Sequencing depths for the variants of results from four companies (AA to DD). The read counts for the variants from Dragen system for each company were plotted. X-axis, chromosome sites for variants. Y-axis, box plot of read counts for variants with the median, top 25%, and lower 75% read counts.

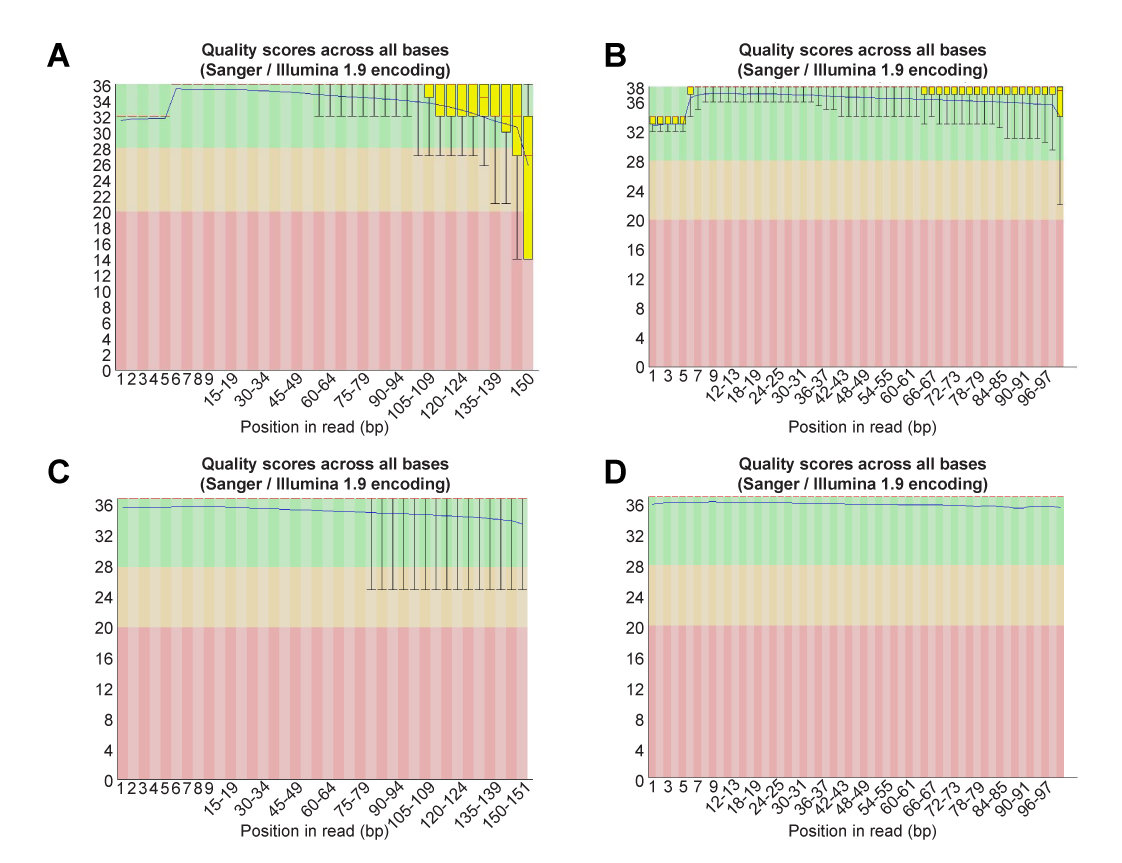

**Fig. S5** Per base sequence quality for raw T-NGS data of DNA2 sample from each company. The results were chosen as representative results for each company. A. Company AA. B. Company BB, C. Company CC. D. Company DD.

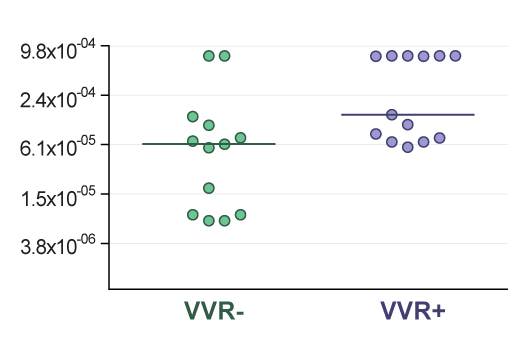

**Fig. S6** Total false positive (FP) error rates depending on the inclusion or exclusion of the VVR errors (V-V pair errors incurred by the appearance of reference bases in V-V pairs) for the T-NGS variant call results from BB company by inhouse method. With the exclusion of VVR errors, the median total FP error rate decreased to about a half. In x-axis, VVR-, total FP error rates with the exclusion of the VVR errors; VVR+, total FP error rates with the exclusion of the VVR errors. The total FP error rate for each circle is from various DNA mixtures between DNA1 and DNA2.

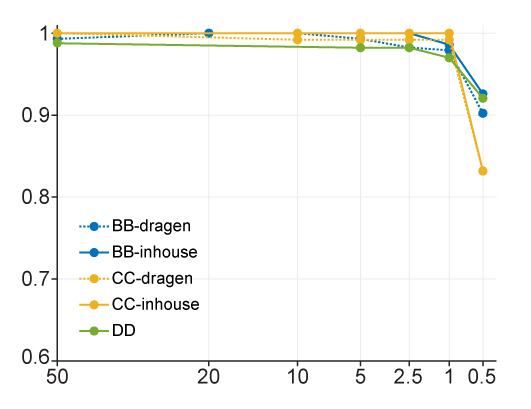

**Fig. S7** Detection rate of diluted reference bases in the mixed DNA reference standards. X-axis, percentage of reference bases in expected allelic fraction. Y-axis, detection rate of reference bases at the informative alleles in reference-standard mixed DNAs. AA-inhouse, T-NGS raw data from AA, and variant call by AA inhouse method; AA-dragen, T-NGS raw data from AA, and variant call by Dragen system; BB-dragen, T-NGS raw data from BB, and variant call by Dragen system; BB-inhouse, T-NGS raw data from BB, and variant call by BB inhouse method; CC-dragen, T-NGS raw data from CC, and variant call by Dragen system; CC-inhouse, T-NGS raw data from CC, and variant call by CC inhouse method; DD, T-NGS raw data from DD, and variant call by DD inhouse method.

**Table S1**. Number of variants in control sample depending on the analytic conditions of default or solid conditions by Dragen system

| Company | Default | Solid | Ratio (%, solid/default) |
| --- | --- | --- | --- |
| AA | 2393 | 140538 | 58.7 |
| BB | 1149 | 23172 | 20.2 |
| CC | 481 | 45886 | 95.4 |

**Table S2**. Basic information of T-NGS kits from 4 companies.

| Company | DNA input | Target size (Mb) | Target gene number |
| --- | --- | --- | --- |
| AA | 200 ng | 3.68 (exon) | 170 |
| BB | 200 ng | 1.66 (exon and promoter) | 335 |
| CC | 200 ng | 7.32 (exon) | 407 |
| DD | 30 ng | 9.54 (exon and intron)) | 523 |

**Table S3**. Target gene list of T-NGS kits from 4 companies

| Company | Target genes |
| --- | --- |
| AA | ABL1, ABL2, AKT1, AKT2, AKT3, ALK, APC, AR, ARAF, ASXL1, ATM, ATR, AURKA, AURKB, AURKC, AXL, BAP1, BCL2, BRAF, BRCA1, BRCA2, BRD2, BRD3, BRD4, CBFB, CCND1, CCND2, CCND3, CCNE1, CDH1, CDK12, CDK4, CDK6, CDKN1A, CDKN1B, CDKN2A, CDKN2B, CDKN2C, CEBPA, CHEK2, CREBBP, CRKL, CSF1R, CTNNB1, DDR1, DDR2, DNMT3A, DOT1L, EGFR, EPHA3, ERBB2, ERBB3, ERBB4, ERCC2, ERG, ERRFI1, ESR1, ETV1, ETV4, ETV5, ETV6, EWSR1, EZH2, FBXW7, FGFR1, FGFR2, FGFR3, FGFR4, FLCN, FLT1, FLT3, FLT4, FOXL2, GNA11, GNAQ, GNAS, HDAC9, HGF, HRAS, IDH1, IDH2, IGF1R, IGF2, JAK1, JAK2, JAK3, KDR, KIT, KMT2A, KRAS, MAP2K1, MAP2K2, MAP2K4, MAP3K1, MAP3K4, MAPK1, MAPK3, MAPK8, MCL1, MDM2, MDM4, MED12, MEN1, MET, MITF, MLH1, MPL, MSH2, MSH6, MTOR, MYC, MYCN, MYD88, NF1, NF2, NFKBIA, NKX2-1, NOTCH1, NOTCH2, NOTCH3, NOTCH4, NPM1, NRAS, NTRK1, NTRK2, NTRK3, NUTM1, PDGFB, PDGFRA, PDGFRB, PIK3CA, PIK3CB, PIK3CD, PIK3R1, PIK3R2, POLE, PPARG, PTCH1, PTEN, RAB35, RAD50, RAF1, RARA, RB1, RET, RHEB, RICTOR, RNF43, ROS1, RSPO1, RSPO2, RUNX1, SMAD2, SMAD4, SMARCA4, SMARCB1, SMO, SRC, STK11, SYK, TET2, TMPRSS2, TOP2A, TP53, TSC1, TSC2, VHL, WT1, XPO1, ZNRF3 |
| BB | ABL1, ABL2, ACVR1B, AKT1, AKT2, AKT3, ALK, ALOX12B, AMER1, APC, AR, ARAF, ARFRP1, ARID1A, ARID1B, ARID2, ASXL1, ATM, ATR, ATRX, AURKA, AURKB, AXIN1, AXL, BAP1, BARD1, BCL2, BCL2L1, BCL2L2, BCL6, BCOR, BCORL1, BLM, BRAF, BRCA1, BRCA2, BRD4, BRIP1, BTG1, BTG2, BTK, C11ORF30, CALR, CARD11, CASP8, CBFB, CBL, CCND1, CCND2, CCND3, CCNE1, CD22, CD274, CD70, CD79A, CD79B, CDC73, CDH1, CDK12, CDK4, CDK6, CDK8, CDKN1A, CDKN1B, CDKN2A, CDKN2B, CDKN2C, CEBPA, CHD2, CHD4, CHEK1, CHEK2, CIC, CREBBP, CRKL, CRLF2, CSF1R, CSF3R, CTCF, CTNNA1, CTNNB1, CUL3, CUL4A, CXCR4, CYLD, CYP17A1, DAXX, DDR1, DDR2, DICER1, DIS3, DNMT1, DNMT3A, DOT1L, EED, EGFR, EP300, EPHA3, EPHA5, EPHA7, EPHB1, EPHB4, ERBB2, ERBB3, ERBB4, ERCC4, ERG, ERRFI1, ESR1, ETV1, ETV6, EZH2, FAM46C, FANCA, FANCC, FANCD2, FANCE, FANCF, FANCG, FANCL, FAS, FAT1, FBXW7, FGF10, FGF12, FGF14, FGF19, FGF23, FGF3, FGF4, FGF6, FGFR1, FGFR2, FGFR3, FGFR4, FH, FLCN, FLT1, FLT3, FLT4, FOXL2, FOXP1, FRS2, FUBP1, GABRA6, GATA1, GATA2, GATA3, GATA4, GATA6, GID4, GLI1, GNA11, GNA13, GNAQ, GNAS, GPR124, GRIN2A, GRM3, GSK3B, H3F3A, HDAC1, HGF, HIST1H3B, HNF1A, HRAS, HSD3B1, HSP90AA1, ID3, IDH1, IDH2, IGF1R, IGF2, IKBKE, IKZF1, IL7R, INHBA, INPP4B, IRF2, IRF4, IRS2, JAK1, JAK2, JAK3, JUN, KAT6A, KDM5A, KDM5C, KDM6A, KDR, KEAP1, KEL, KIT, KLHL6, KMT2A, KMT2C, KMT2D, KRAS, LMO1, LRP1B, LTK, LYN, LZTR1, MAF, MAGI2, MAP2K1, MAP2K2, MAP2K4, MAP3K1, MAP3K13, MAPK1, MBIP, MCL1, MDM2, MDM4, MED12, MEF2B, MEN1, MERTK, MET, MITF, MKNK1, MLH1, MPL, MRE11A, MSH2, MSH3, MSH6, MSI-01, MSI-03, MSI-06, MSI-07, MSI-08, MSI-11, MSI-13, MSI-14, MST1R, MTAP, MTOR, MUTYH, MYC, MYCL, MYCN, MYD88, NBN, NF1, NF2, NFE2L2, NFKBIA, NKX2-1, NOTCH1, NOTCH2, NOTCH3, NPM1, NRAS, NSD1, NT5C2, NTRK1, NTRK2, NTRK3, NUP93, P2RY8, PAK3, PAK7, PALB2, PARK2, PARP1, PARP2, PARP3, PAX5, PBRM1, PDCD1, PDCD1LG2, PDGFRA, PDGFRB, PDK1, PIK3C2B, PIK3C2G, PIK3CA, PIK3CB, PIK3CG, PIK3R1, PIK3R2, PIM1, PLCG2, PMS2, POLD1, POLE, PPARG, PPP2R1A, PPP2R2A, PRDM1, PREX2, PRKAR1A, PRKCI, PRKDC, PRSS8, PTCH1, PTEN, PTPN11, PTPRD, PTPRO, QKI, RAC1, RAD21, RAD50, RAD51, RAD51B, RAD51C, RAD51D, RAD52, RAD54L, RAF1, RANBP2, RARA, RB1, RBM10, REL, RET, RHOA, RICTOR, RIT1, RNF43, ROS1, RPTOR, RUNX1, RUNX1T1, SDHA, SDHB, SDHC, SDHD, SETD2, SF3B1, SGK1, SLIT2, SMAD2, SMAD3, SMAD4, SMARCA4, SMARCB1, SMO, SNCAIP, SOCS1, SOX10, SOX2, SOX9, SPEN, SPOP, SPTA1, SRC, STAG2, STAT3, STAT4, STK11, SUFU, SYK, TAF1, TBX3, TEK, TERC, TERT, TET2, TGFBR2, TIPARP, TNFAIP3, TNFRSF14, TOP1, TOP2A, TP53, TSC1, TSC2, TSHR, TYRO3, U2AF1, VEGFA, VHL, WHSC1, WHSC1L1, WISP3, WT1, XPO1, XRCC2, ZBTB2, ZNF217, ZNF703 |
| CC | MET, TERT, ALK, RET, ROS1, TMPRSS2, ABCB1, ABCC1, ABCC3, ABCG2, ABL1, ABL2, AKT1, AKT2, AKT3, APC, APCDD1, APLNR, APOBEC3A, APOBEC3B, AR, ARAF, AREG, ARID1A, ARID1B, ARID2, ASNS, ASXL1, ATM, ATR, ATRX, AURKA, AURKB, AXIN1, AXIN2, AXL, B2M, BACH1, BAP1, BARD1, BCL2, BCL2A1, BCL2L11, BCL2L2, BCL6, BCOR, BCORL1, BIRC7, BLM, BRAF, BRCA1, BRCA2, BRD7, BRIP1, CALR, CARD11, CASP8, CBFB, CBL, CBLB, CCND1, CCND2, CCND3, CCNE1, CD274, CD79A, CD79B, CDC73, CDH1, CDH2, CDH5, CDK1, CDK12, CDK4, CDK6, CDK8, CDKN1A, CDKN1B, CDKN2A, CDKN2B, CDKN2C, CDX2, CEBPA, CHD4, CHEK1, CHEK2, CHUK, CIC, CREBBP, CSF1R, CSF3R, CTNNB1, CUL4A, CYP17A1, DDR2, DICER1, DIS3, DNMT1, DNMT3A, DOT1L, DPYD, EGF, EGFR, EIF1AX, EML4, EP300, EPHA2, EPHA3, EPHA6, EPHA7, EPHB1, EPHB4, EPHB6, ERBB2, ERBB3, ERBB4, ERCC1, ERCC2, ERCC3, ERCC4, ERCC5, ERCC6, ERG, ESR1, ETS1, ETV1, ETV4, ETV5, ETV6, EWSR1, EZH2, FAM175A, FANCA, FANCC, FANCD2, FANCE, FANCF, FANCG, FANCI, FANCL, FANCM, FAT1, FAT3, FBXW7, FCGR2B, FGF1, FGF10, FGF14, FGF18, FGF19, FGF2, FGF23, FGF3, FGF4, FGF5, FGF6, FGF7, FGF8, FGF9, FGFR1, FGFR2, FGFR3, FGFR4, FLCN, FLI1, FLT1, FLT3, FLT4, FOXA1, FOXL2, FOXP1, FRS2, G6PD, GATA1, GATA2, GATA3, GEN1, GLI2, GNA11, GNAQ, GNAS, GSTP1, H19, H3F3A, HDAC1, HDAC2, HELQ, HGF, HIF1A, HIST1H3B, HLA-A, HLA-B, HLA-C, HLA-DPA1, HLA-DPB1, HLA-DQA1, HLA-DQB1, HLA-DRA, HLA-DRB1, HLA-DRB5, HNF1A, HRAS, HSP90AA1, HSPH1, IDH1, IDH2, IDO1, IGF1, IGF1R, IGF2, IGF2R, IKBKE, IKZF1, IL1B, IL7R, INPP4B, INSR, IRF1, IRS2, ITK, JAK1, JAK2, JAK3, KAT6A, KCNJ5, KDM5A, KDR, KIF5B, KIT, KMT2A, KMT2D, KRAS, LAMP1, LATS1, LATS2, LRP1B, LRP6, LTK, MAD1L1, MAP2K1, MAP2K2, MAP2K4, MAP2K7, MAP3K1, MAP3K13, MAPK1, MCL1, MDC1, MDM2, MDM4, MED12, MEN1, MITF, MLH1, MLLT3, MPL, MRE11A, MSH2, MSH3, MSH6, MTHFR, MTOR, MUS81, MUTYH, MYC, MYCL1, MYCN, MYD88, NBN, NCOA3, NCOR1, NF1, NF2, NFKBIA, NKX2-1, NOTCH1, NOTCH2, NOTCH3, NOTCH4, NPM1, NRAS, NRG1, NSD1, NT5C2, NTRK1, NTRK2, NTRK3, PAK1, PAK7, PALB2, PARP1, PARP2, PARP3, PARP4, PAX3, PAX7, PBRM1, PDCD1LG2, PDGFRA, PDGFRB, PDPK1, PIK3CA, PIK3CB, PIK3CD, PIK3CG, PIK3R1, PIK3R2, PKHD1, PLCG1, PML, PMS2, POLD1, POLE, POLQ, PPARG, PPP1R15A, PPP2R2A, PRDM1, PRKCB, PTCH1, PTCH2, PTEN, PTPN11, PTPRD, RAB35, RAC1, RAD21, RAD50, RAD51, RAD51B, RAD51C, RAD51D, RAD54L, RAF1, RB1, RELA, RHBDF2, RICTOR, RIT1, RNF43, RPS6KB1, RPTOR, RRAS2, RSF1, RUNX1, SDHAF2, SDHB, SDHC, SDHD, SERPINB3, SETBP1, SETD2, SF3B1, SH2B3, SLC29A1, SLX4, SMAD2, SMAD4, SMARCA1, SMARCA4, SMARCB1, SMO, SOCS1, SOX2, SPEN, SPOP, SRC, SRSF2, STAG2, STAT1, STAT3, STAT4, STK11, SUZ12, SYK, TBX3, TET2, TFRC, TGFBR1, TGFBR2, TNFAIP3, TNFRSF14, TNKS, TOP1, TOP2A, TP53, TP53BP1, TPMT, TSC1, TSHR, TTF1, TYMS, U2AF1, UGT1A1, VEGFA, VHL, WEE1, WRN, WT1, XPA, XPC, XRCC1, XRCC2, XRCC3, ZBTB16, ZNF217, ZNRF3 |
| DD | ABL1, ABL2, ACVR1, ACVR1B, AKT1, AKT2, AKT3, ALK, ALOX12B, ANKRD11, ANKRD26, APC, AR, ARAF, ARFRP1, ARID1A, ARID1B, ARID2, ARID5B, ASXL1, ASXL2, ATM, ATR, ATRX, AURKA, AURKB, AXIN1, AXIN2, AXL, B2M, BAP1, BARD1, BBC3, BCL10, BCL2, BCL2L1, BCL2L11, BCL2L2, BCL6, BCOR, BCORL1, BCR, BIRC3, BLM, BMPR1A, BRAF, BRCA1, BRCA2, BRD4, BRIP1, BTG1, BTK, C11orf30, CALR, CARD11, CASP8, CBFB, CBL, CCND1, CCND2, CCND3, CCNE1, CD274, CD276, CD74, CD79A, CD79B, CDC73, CDH1, CDK12, CDK4, CDK6, CDK8, CDKN1A, CDKN1B, CDKN2A, CDKN2B, CDKN2C, CEBPA, CENPA, CHD2, CHD4, CHEK1, CHEK2, CIC, CREBBP, CRKL, CRLF2, CSF1R, CSF3R, CSNK1A1, CTCF, CTLA4, CTNNA1, CTNNB1, CUL3, CUX1, CXCR4, CYLD, DAXX, DCUN1D1, DDR2, DDX41, DHX15, DICER1, DIS3, DNAJB1, DNMT1, DNMT3A, DNMT3B, DOT1L, E2F3, EED, EGFL7, EGFR, EIF1AX, EIF4A2, EIF4E, EML4, EP300, EPCAM, EPHA3, EPHA5, EPHA7, EPHB1, ERBB2, ERBB3, ERBB4, ERCC1, ERCC2, ERCC3, ERCC4, ERCC5, ERG, ERRFI1, ESR1, ETS1, ETV1, ETV4, ETV5, ETV6, EWSR1, EZH2, FAM123B, FAM175A, FAM46C, FANCA, FANCC, FANCD2, FANCE, FANCF, FANCG, FANCI, FANCL, FAS, FAT1, FBXW7, FGF1, FGF10, FGF14, FGF19, FGF2, FGF23, FGF3, FGF4, FGF5, FGF6, FGF7, FGF8, FGF9, FGFR1, FGFR2, FGFR3, FGFR4, FH, FLCN, FLI1, FLT1, FLT3, FLT4, FOXA1, FOXL2, FOXO1, FOXP1, FRS2, FUBP1, FYN, GABRA6, GATA1, GATA2, GATA3, GATA4, GATA6, GEN1, GID4, GLI1, GNA11, GNA13, GNAQ, GNAS, GPR124, GPS2, GREM1, GRIN2A, GRM3, GSK3B, H3F3A, H3F3B, H3F3C, HGF, HIST1H1C, HIST1H2BD, HIST1H3A, HIST1H3B, HIST1H3C, HIST1H3D, HIST1H3E, HIST1H3F, HIST1H3G, HIST1H3H, HIST1H3I, HIST1H3J, HIST2H3A, HIST2H3C, HIST2H3D, HIST3H3, HLA-A, HLA-B, HLA-C, HNF1A, HNRNPK, HOXB13, HRAS, HSD3B1, HSP90AA1, ICOSLG, ID3, IDH1, IDH2, IFNGR1, IGF1, IGF1R, IGF2, IKBKE, IKZF1, IL10, IL7R, INHA, INHBA, INPP4A, INPP4B, INSR, IRF2, IRF4, IRS1, IRS2, JAK1, JAK2, JAK3, JUN, KAT6A, KDM5A, KDM5C, KDM6A, KDR, KEAP1, KEL, KIF5B, KIT, KLF4, KLHL6, KMT2B, KMT2C, KMT2D, KRAS, LAMP1, LATS1, LATS2, LMO1, LRP1B, LYN, LZTR1, MAGI2, MALT1, MAP2K1, MAP2K2, MAP2K4, MAP3K1, MAP3K13, MAP3K14, MAP3K4, MAPK1, MAPK3, MAX, MCL1, MDC1, MDM2, MDM4, MED12, MEF2B, MEN1, MET, MGA, MITF, MLH1, MLL, MLLT3, MPL, MRE11A, MSH2, MSH3, MSH6, MST1, MST1R, MTOR, MUTYH, MYB, MYC, MYCL1, MYCN, MYD88, MYOD1, NAB2, NBN, NCOA3, NCOR1, NEGR1, NF1, NF2, NFE2L2, NFKBIA, NKX2-1, NKX3-1, NOTCH1, NOTCH2, NOTCH3, NOTCH4, NPM1, NRAS, NRG1, NSD1, NTRK1, NTRK2, NTRK3, NUP93, NUTM1, PAK1, PAK3, PAK7, PALB2, PARK2, PARP1, PAX3, PAX5, PAX7, PAX8, PBRM1, PDCD1, PDCD1LG2, PDGFRA, PDGFRB, PDK1, PDPK1, PGR, PHF6, PHOX2B, PIK3C2B, PIK3C2G, PIK3C3, PIK3CA, PIK3CB, PIK3CD, PIK3CG, PIK3R1, PIK3R2, PIK3R3, PIM1, PLCG2, PLK2, PMAIP1, PMS1, PMS2, PNRC1, POLD1, POLE, PPARG, PPM1D, PPP2R1A, PPP2R2A, PPP6C, PRDM1, PREX2, PRKAR1A, PRKCI, PRKDC, PRSS8, PTCH1, PTEN, PTPN11, PTPRD, PTPRS, PTPRT, QKI, RAB35, RAC1, RAD21, RAD50, RAD51, RAD51B, RAD51C, RAD51D, RAD52, RAD54L, RAF1, RANBP2, RARA, RASA1, RB1, RBM10, RECQL4, REL, RET, RFWD2, RHEB, RHOA, RICTOR, RIT1, RNF43, ROS1, RPS6KA4, RPS6KB1, RPS6KB2, RPTOR, RUNX1, RUNX1T1, RYBP, SDHA, SDHAF2, SDHB, SDHC, SDHD, SETBP1, SETD2, SF3B1, SH2B3, SH2D1A, SHQ1, SLIT2, SLX4, SMAD2, SMAD3, SMAD4, SMARCA4, SMARCB1, SMARCD1, SMC1A, SMC3, SMO, SNCAIP, SOCS1, SOX10, SOX17, SOX2, SOX9, SPEN, SPOP, SPTA1, SRC, SRSF2, STAG1, STAG2, STAT3, STAT4, STAT5A, STAT5B, STK11, STK40, SUFU, SUZ12, SYK, TAF1, TBX3, TCEB1, TCF3, TCF7L2, TERC, TERT, TET1, TET2, TFE3, TFRC, TGFBR1, TGFBR2, TMEM127, TMPRSS2, TNFAIP3, TNFRSF14, TOP1, TOP2A, TP53, TP63, TRAF2, TRAF7, TSC1, TSC2, TSHR, U2AF1, VEGFA, VHL, VTCN1, WISP3, WT1, XIAP, XPO1, XRCC2, YAP1, YES1, ZBTB2, ZBTB7A, ZFHX3, ZNF217, ZNF703, ZRSR2 |
